# Elucidating the Molecular Basis of pH Activation of an Engineered Mechanosensitive Channel

**DOI:** 10.1101/707794

**Authors:** Kalyan Immadisetty, Adithya Polasa, Reid Shelton, Mahmoud Moradi

## Abstract

Mechanosensitive (MS) channels detect and respond to changes in the pressure profile of cellular membranes and transduce the mechanical energy into electrical and/or chemical signals. However, by re-engineering the MS channels, chemical signals such as pH change can trigger the activation of some MS channels. This paper elucidate the activation mechanism of an engineered MS channel of large conductance (MscL) at an atomic level through a combination of equilibrium, non-equilibrium, biased, and unbiased molecular dynamics (MD) simulations for the first time. Comparing the wild-type and engineered MscL activation processes at an atomic level suggests that the two systems are likely to be associated with different active states and different transition pathways. These findings indicate that (1) periplasmic loops play a key role in the activation process of MscL, (2) the loss of various hydrogen bonds and salt bridge interactions in the engineered MscL channel causes the spontaneous opening of the channel, and (3) the most significant interactions lost during the activation process are those between the transmembrane (TM) helices 1 and 2 (TM1 and TM2) in engineered MscL channel. In this research, the orientation-based biasing approach for producing and optimizing an open MscL model is a promising way to characterize unknown protein functional states and to research the activation processes in ion channels. String method with swarms of trajectories (SMwST) was used to identify the optimal transition pathway and elucidate the activation mechanism of the engineered MscL. Finally, the free energy profile of engineered MscL associated with the activation process using a novel along-the-path free energy calculation approach is constructed. This work paves the way for a computational framework for the studies aimed at designing pH-triggered channel-functionalized drug delivery liposomes.

## Introduction

In bacteria, MscL channels serve as an emergency release valve for acute osmolarity in the system, preserving osmotic homeostasis in bacterial cells. ^2^ MscL has been a prominent drug target for many antibiotics^3,4^ due to its direct interaction with bacterial potency. ^5^

MscL is necessary in the growth of bacteria and any mutation in the protein could cause severe modifications.^4,6^

MscL protein is specifically found in many bacterial cell members but absent in mammalian genomes. ^4^,

In mammalian cells, engineered MscL channels could be expressed genetically.^7^ MscL can be used in absorption of membrane-impermeable drugs and bioactive materials into cells using ultrasonic^8^ or chemical ^7^ activation processes. Introduction of bacterial MscL channel protein in mice has shown a reduced metastasis in lungs.^9^ However, engineered MscL expressed in rat hippocampal neurons is activated in low pressure ultrasound pulse which controls neuronal excitation. ^8^

MscL is an rv80-kDa homo-pentameric membrane protein with each pr otomer consisting of two *α*-helical transmembrane (TM) helices. ^2, 10^ MscL channels identify and react to changes in the pressure profile of cell membrane and transduce the mechanical energy into electrical or potentially chemical signals. ^11–14^ However, by re-designing, the activation of MscL can be set off by chemical signals. For example, pH change, ^15–22^ which is a basis for using an engineered MscL as a pH-sensing nanovalve in a drug delivery liposome (DDL) ^10, 16, 23^ (Fig. 1). Targeted modification of cystine containing subunits with positively charged thiol groups have determined to be functional when inserted in liposomes. ^24^ In G22C mutated system, MscL is modeled with [2-(trimethylammonium)ethyl] methane thiosulfonate bromide (MTSET) which resulted in spontaneous opening of the channel even in absence of natural trigger and in physiological conditions. ^10, 20, 22^ Varying the p*K_a_* and hydrophobicity of the modulators could result in the activation of pH-induced opening of the channel at different desired pH conditions. ^16, 17, 23^

**Fig. 1.**
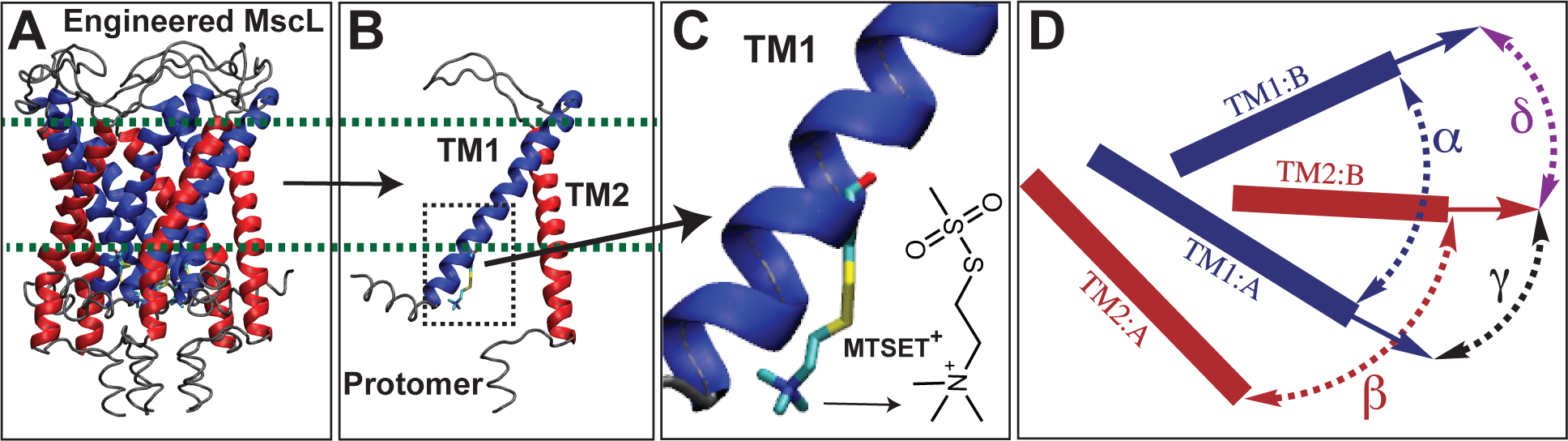
Engineered MscL channel. (A) Cartoon representation of engineered tb-MscL model based on the crystal structure of wild-type tb-Msc1.^1^ (B) A protomer of engineered tb-MscL. (C) Transmembrane helix 1 (TMl) of engineered tb-MscL. pH-sensitive labels at­ tached to A20C tbMscL mutant enable a pH-dependent activation mechanis m for MscL channel. [2-(trimethylammonium)ethyl] methane thiosulfonate bromide (MTSET+) attach­ ment activates MscL even in physiological conditions. (D) Schematic representation of angles calculated for analysis, *a* is the angle between TMls of neighboring protomers, *f3* is the angle between TM2s of the neighboring protomers, 1is the angle between TMl and TM2 of the neighboring protomers, and *5* is the angle between TMl and TM2 of same domain.

Various molecular dynamics (MD) studies have been conducted to study the wild-type MscL activation, often involving coarse-graining ^25–27^ or conventional biased approaches. ^28–41^

Other computational approaches such as static molecular modeling ^42^ and continuum models ^43–45^ have been used for the study of wild-type MscL activation as well. For the first time, the activation mechanism of engineered MscL by pH changes at an atomic level and within the thermodynamic conditions was investigated. Also, an all-atom equilibrium un-biased MD simulations followed by non-equilibrium simulations are conducted, in which system-specic collective variables have been used to apply force and impose rotational changes on MscL TM helices using the orientation quaternion technique. ^46–50^ These simulations are then followed by string method with swarms of trajectories and free energy calculations to fully describe the activation process of engineered MscL.

Observation in the initial unbiased equilibrium MD indicated that the introduction of MTSET labels partially open the channel, especially near the narrowest region of the pore in the intracellular side. However, the channel does not open completely on the extracellular side within the timescale of simulations (1 microsecond). Non-equilibrium simulations per-formed to determine the maximum extent to which the channel opens. Moreover, the open state structures of the channel resulting from non-equilibrium simulations are relaxed via un-biased equilibrium simulations in order to examine whether the structures are stable in open state. The final conformation of non-equilibrium wild-type (WT) structure collapsed back to the closed conformation in the follow up equilibrium simulations, while the engineered MscL structure stably remained in open conformational state in follow up equilibrium state.

In more detail, engineered MscL activation is analyzed by relaxing the transition path using a string method with swarms of trajectory (SMwST), followed by a novel method of free energy estimation based on unbiased MD simulations along the minimum free energy pathway. Finally, based on the performed extensive simulations, a mechanism for the activation of the channels proposed. Overall, our computational results are in line with and may better explain different lab studies that could not be completely understood due to the limitations of experimental techniques. For instance, the engineered MscL shows a higher dynamic activity compared to the WT.

The effect of different simulation conditions on MscL’s conformational dynamics within time scales accessible to normal MD simulations is explored in this detailed research review. To this effect, all-atom MD simulations of MscL are conducted comprehensively. In terms of WT and engineered, 10 separate simulations of MscL under different conditions were carried out. A detailed study of our MD simulation trajectories shows a separate behavior of simulation conditions of local conformational dynamics. In the following, the simulation specifics in theoretical methods are discussed first. An description of the measurements and quantities used for the study of our trajectories will be given by the findings and discussion, accompanied by a more detailed account of the results, explaining both local and global conformational fluctuations. For a short description of our observations, the last segment is retained for conclusions.

## Methods

Crystal structure of MscL in closed (in-active) state (PDB: 2OAR) ^1^ was downloaded from pdb.org. Initially the system was prepared using the Molecular Operating Environment (MOE) software ^51^ by removing the crystal waters, assigning the appropriate protonation states for the residues using protonate3D facility and also by adding hydrogens and other missing atoms. Further, CHARMM-GUI web-server ^52, 53^ was used to place the protein in the 1-palmitoyl-2-oleoyl-sn-glycero-3-phosphocholine (POPC) membrane bilayer and also to solvate and build the system. 0.15 mM NaCl was added to the system, generating a total of 179 sodium and 184 chloride ions. Overall, the total size of the system was ≈150×150×130 Å^3^.

The total number of lipids were 603 (298 lipids in the upper leaflet and 305 lipids in the lower leaflet) and the system contained 60,027 TIP3P ^54^ water molecules. The total number of atoms in the system were approximately 270,524. NAMD2.10/2.13 ^55^ was used to simulate the system in periodic boundary conditions in the NPT ensemble at 310 K using a Langevin integrator with a damping coefficient of *γ* =0.5 ps*^−^*^1^ and 1 atm pressure was maintained using the Nośe-Hoover Langevin piston method. ^56, 57^ Time step used was 2 fs and the system was equilibrated for 1000 ns.

CHARMM36 all-atom additive force field parameters were used to simulate the entire system. ^58, 59^ Prior to equilibration, each parent system (Set1) was first energy minimized for 10,000 steps via conjugate gradient algorithm. ^60^ Further, The parent structures were also relaxed using a sequence of *∼*1 ns ^52^ multi-step restraining simulations in an NVT ensemble. The non-bonded interactions were cutoff at 10*−*12 Å and the long-range electrostatic interactions were computed with particle mesh Ewald (PME) method. ^61^ A second system was also generated by mutating A20 in each MscL subunit to cysteine. Next, a pH-sensing label MT-SET was attached to each mutated residue (total 5 labels). Further, the system was prepared using the procedure explained above. The size of the system and the total number of atoms were approximately similar to the above, except that this system has extra labels attached to the cysteines at position 20. CHARMM General Force Field (CGenFF) ^62^ parameters was used for the MTSET labels. A third system was also prepared to serve as a control. In this system, A20 in only one subunit was mutated to cysteine and pH-sensing label MTSET was attached. System was prepared using the similar protocol explained above. Systems 2 and 3 were also simulated each for 1000 ns. Two additional simulations repeats (sets 2 & 3) were also conducted for each of the three systems listed above. The two additional simulations were continued from the first simulations by using the configurations at 150ns and 200ns as the input in all three cases. Since the labels attached to monomers 2 & 4 in 5-MTSET were oriented differently from the other 3, they were manually re-oriented similar to the other 3 and called this 5-MTSET(*) (Fig. S1). One data point for 1 ns was collected for the statistical analysis. Trajectories were visualized using VMD. ^63^ Principle component analysis (PCA) was carried using PRODY software ^64^ on an ensemble of DCD structures and 20 modes were generated. Only C*α* atoms were considered for the PCA calculations. The details of the methodology can be found else where. ^65, 66^ Hydrogen bond and salt bridge interaction analysis was conducted via VMD plugins. The cut-off distance and angle were 3.5Å and 30*^◦^* respectively, for the hydrogen bond analysis. Only one hydrogen bond for an interaction pair was counted. For salt bridge analysis, the cut-off distance was 4 Å and the distance between the oxygen atoms of the acidic residues and nitrogens of basic residues was calculated.

Many mechanically suitable system-specific reaction-coordinates have been established in this research. In this article, the terms reaction coordinates and collective variables are used synonymously. The methodology offers a basic framework for optimizing biasing protocols in a sequence of brief simulations. One can significantly boost the performance of search protocols in sampling complex transformation paths by using advanced system-specific biasing protocols. In the non-equilibrium simulations, equilibrated structures WT, 5-MTSET and 5-MTSET(*) have been opened in an 100ns simulation each using orientation quaternions of different helices, bundles of helices or their linear combinations. A set of orientations were used including Q*_i_* in which Q*_i_* are orientation quaternions of i*^t^h* TM helix. ^50, 67^ Quaternion Q=(q0, q1, q2, q3) can be regarded as a composite of a scalar q0 and an ordinary vector q=(q1, q2, q3), or a complex number Q = q0+q1i+q2j+q3k with three distinct imaginary parts. A quaternion unit will define the perfect rotation to overlay one set of co-ordinates, that is, an orientation quaternion in which a unit is an optimal angle and axis of rotation, respectively. ^50, 67^

For the non-equilibrium simulations, a single molecule FRET (smFRET) based open state model of Mscl ^68^ was used as a target. Each simulation was repeated twice and the force constant used was 10,000 *kcal/mol.rad*^2^ (Table 2). The open state model was equilibriated for 260ns without any constraints in order to validate its conformational consistency.

**Table 1:** Equilibrium MD simulations.

|  | WT | 5-MTSET | 1-MTSET | 5-MTSET(*) |
| --- | --- | --- | --- | --- |
| Set-1 | 1000ns | 1000ns | 1000ns | 500ns |
| Set-2 | 500ns | 500ns | 500ns |  |
| Set-3 | 500ns | 500ns | 500ns |  |

**Table 2:** Protocol for non-equilibrium and the follow up equilibrium simulations.

| Step | Force Constant (k)<br>($kcal/mol.rad^2$) | Time<br>(ns) |
| --- | --- | --- |
| 1 | 10,000 | 100 |
| 2 | 10,000 | 5 |
| 3 | 10,000 to 1000 | 5 |
| 4 | 1000 to 0 | 10 |
| 5 | 0 | 260 |

A parallel version of the string method with the swarms trajectory (SMwST) ^69^ technique was implemented here, in which each image consists of multiple parallel copies, constrained and released iteratively with a periodic exchange of data between the replicas to update the image centers at the end of each iteration. This initial pathway have been extracted from the NE simulation. The engineered MscL activation pathway obtained from one of the non-equilibrium simulations and the 260 ns follow-up equilibrium simulation were relaxed in the string method simulations. The SMwST algorithm begins from an initial string specified by N points/images *xi*, where i is any integer between 1 and N. Colvar *ζ* mainly determines biasing potential, which is U*_i_* = k/2 (*ξ*-*ξ*_0_)^2^ for M copies of image i. From the initial series, initial values for image centers are determined: *ζ* I=*ζ* (xi). The SMwST algorithm consists of three iterative steps. (Step I) Restraining: each device is constrained to use the harmonic potential mentioned above based on the current picture i. (Step II) Drifting: Simulations are started after release from restraints for D (drifting). (Step III) Reparametrization: The new center for each picture I is calculated by combining all observed M-system values associated with picture i at the time of picture i and using a linear interpolation algorithm to hold image centers equidistant. The string converges over these steps to the zero-drift direction, around which the string centers oscillate. 180 iterations of string method were performed to reach a reasonable convergence and the number of copies per image used was 20, for 50 images were used, totalling to 1000 replicas. Each iteration included 5 ps of restraining and 5 ps of unbiased (drifting) simulations. The collective variables included 10 orientation collective variables as discussed above. The restraining force constant was 10,000 *kcal/mol.rad*^2^ (Table 3). Following the SMwST simulation, unbiased MD simulations are conducted for ∼2 ns, starting with the last snapshot of each image copy (1000 simulations) of SMwST simulations. Then post SMwST MD trajectories are used to build an empirical transition matrices ^70^ using a given lag time. Using a novel approach developed by Goolsby *et .al.,* ^70^ this transition matrices were used to estimate both the free energy profile and the activation rate.

**Table 3:** Protocol for string method simulations.

| Step | Force Constant (k)<br>($kcal/mol.rad^2$) | images | copies/image | iterations |
| --- | --- | --- | --- | --- |
| 1 | 10,000 | 50 | 20 | 180 |
| 2 | 0 | 50 | 20 |  |

## Results and Discussion

### MscL spontaneously responds to the attachment of MTSET labels by starting to open

First, to examine the stability of the systems, the protein *C_α_* RMSD (Fig. 2) was measured as a function of time during various equilibrium simulations. RMSD of the WT MscL, in all three sets remained under 5 Å (Fig. 2A). In the case of 1-MTSET and 5-MTSET, RMSDs went up within the first 200ns and remained within 6-8 Å (see Fig. 2B for 5-MTSET and Fig. 2C for 1-MTSET). This highlights the fact that the impact of labels is spontaneous. This observation also establishes that even one label may trigger enough disturbance in the protein environment to trigger an opening, although the protein response is greater when there are 5 labels rather than one. RMSD of the entire protein follows the same pattern as RMSD of extracellular loops (Figs. S3 A, B, and C), highlighting the fact that the loops are driving or dominating the changes in the protein. This is quiet interesting keeping in mind the fact that although labels are attached at the intracellular side, the immediate impact is seen not just on the intracellular side, but also on the extracellular side, which is evidenced by the immediate rise in RMSD of the loops.

**Fig. 2.**
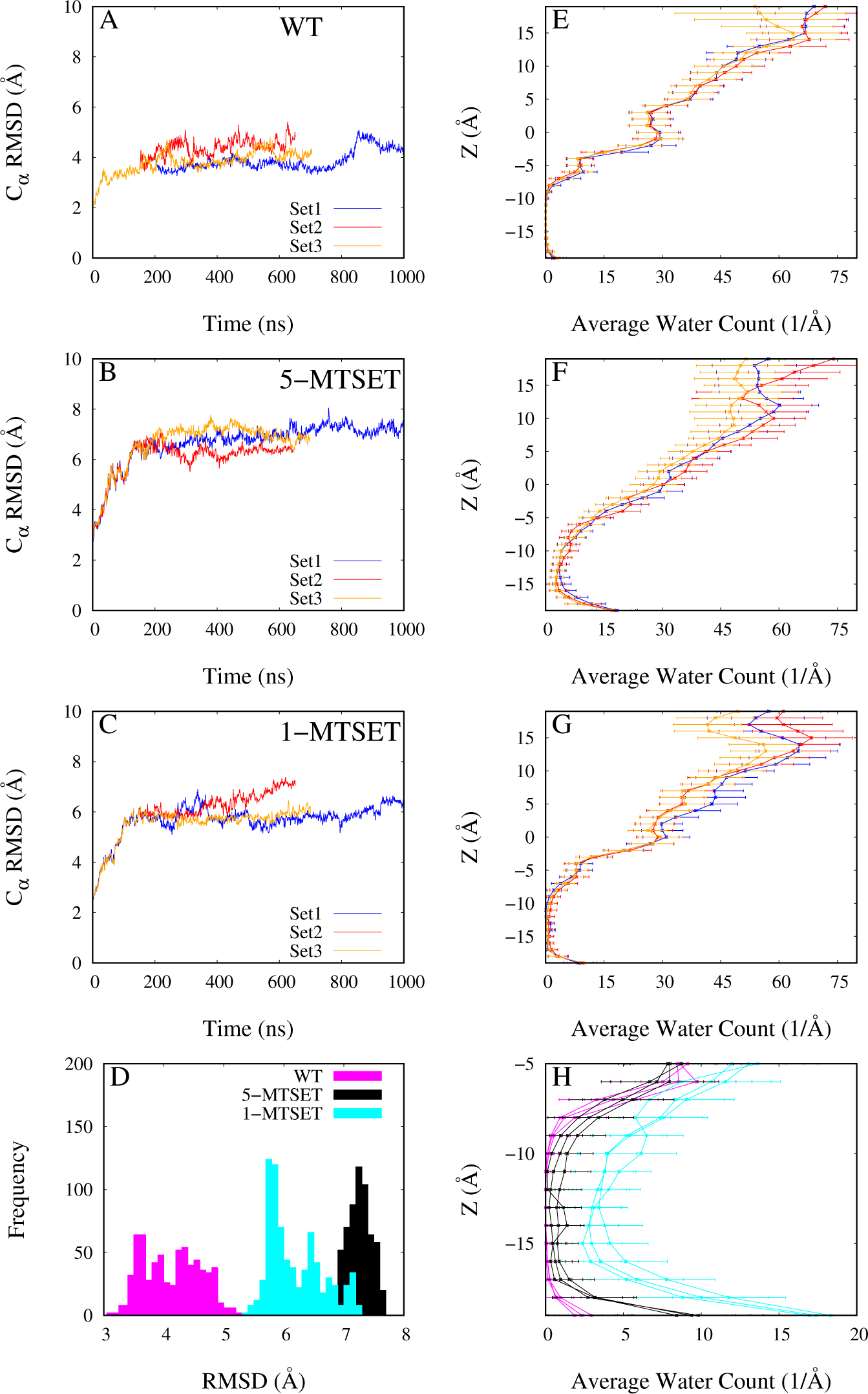
Protein RMSD (Ca.) and water count across the channel pore. (A-C) WT (A) RMSD was stablized around 4 A and the 5-MTSET (B) were distributed between 6-8 A, while the 1-MTSET was stabillized around 6A, except the set-2 simulation (C). (D) Frequency distribution of these RMSDs were shown in D. Only the last 300ns of the trajectory was considered for generating the frequency plot. (E-H) Average water count with respective to Z axis of channel pore. Z between -9 and -19 A reflects the narrower region of the channel that restricts the passage of the ligands across the channel.

To see the effect of labeling on specific area of the protein, the mean RMSF of the C*_α_* atoms of each residue are measured. Irrespective of the system, extracellular loops are dominating compared to the rest of the protein, although, the extent of fluctuation varies between the systems. RMSF of the entire protein in the case of WT is between 0.5-6 Å (Fig. S2A), for 1-MTSET 0.5-7 Å except for first sub-unit 1 (S1), where it is fluctuating between 0.5-11 Å (Fig. S2B). For the 5-MTSET, RMSF of all sub-units are evenly fluctuating between 0.5-7 Å except for sub-unit 2 (S2), where it went upto, 9.5 Å (Fig. S2C). RMSF of TM helices in WT are varying between 0.5-2 Å (Fig. S2D), where as in 1-MTSET between 0.7-3 Å (Fig. S2E), and in other cases between 1.0-3.5 Å (Fig. S2F). This shows that the impact of the labels in the 5-MTSET systems is symmetrical on the TM helices, although it varies in the extracellular loops.

### MTSET labels trigger a partial spontaneous opening of the channel

To estimate the extent of opening of the channel, the water content across the pore was calculated. Throughout all sets of equilibrium simulations, the WT MscL remains completely closed, and there are no water molecules around the intracellular gate region (Z = -10 to -17 Å, Fig. 2E). In the case of 1-MTSET, single label failed to open the channel, although there are some water molecules in the gate region (Fig. 2G). In the case of 5-MTSET (sets 1, 2 and 3), the channel is more open compared to the WT and 1-MTSET (Fig. 2F). Fig. 2H specifically compares the water content around the intracellular gate region (Fig. 2B). Overall the extent of the opening of the channel, particularly near the intracellular gate region is as follows 5-MTSET *>* 1-MTSET *>* WT (Fig. 2H). Apart from the intracellular gate there were several different behavior towards the extracellular opening of the pore (Z = 15-20 Å). While the extent of opening in this region is somewhat reversed as compared to the intracellular gate (i.e., WT *>* 5-MTSET *>* 1-MTSET Figs. 2 E, F, and G). This is due to the collapse of the extracellular loops into the center of the channel pore in the case of 1-MTSET and 5-MTSET and blocking the pore on the extracellular side, which does not happen in the WT (Figs. S3 C, D, E, and F). Overall, based on our water content analysis one could conclude that 5-MTSET is more effective than 1-MTSET in opening the channel. Hence, 1-MTSET system is not considered for further analysis in the rest of the paper.

### MTSET labels disrupt the hydrogen bond network in the MscL

A detailed hydrogen bond (H-Bond) interaction analysis was carried out and discovered that the H-Bond interaction pattern is disturbed by MTSET labels in the MscL channel, particularly backbone-backbone (BB-BB) H-Bonds. The total number of unique BB-BB H-Bond frequency of each residue pair in the WT is greater than in the 5-MTSET (≈5%), and this trend is seen in the entire trajectory (Figs. 3 A, B, and C). The trend was also reflected in TM1 and TM2. This has contributed to the conclusion that the loss of BB-BB H-Bonds in the engineered proteins makes the TM helices more flexible, which is favorable for the conformational changes that are expected to happen.

Two significant inter-helical side chain-side chain (SC-SC) H-Bond interactions were identified with a more significant interaction frequency for WT as compared to the 5-MTSET including the N70-N44 and D16-Y94 residue pairs. N70-N44 is formed between the TM1 and TM2 of the same protomer (denoted by (i,i)) on the extracellular side (Fig. 3H). D16-Y94 is formed between the TM1 (D16) and TM2 (Y94) of neighboring protomers (denoted by (i,i+4), where TM1 of i’th protomer interacts with the TM2 of (i+4)’th protomer) on the intracellular side (Fig. 3H). The average interaction frequency of N70-N44 in the WT is ≈ 50, 40, and 52%, and that of the 5-MTSET is ≈ 20, 22, and 39% for sets 1, 2 and 3, respectively (Fig. 3F), whereas that of the D16-Y94 in WT and 5-MTSET are *≈* 49, 49, and 40% and *≈* 10, 21, and 19%, respectively (Fig. 3G). The N70-N44 interaction might play a key role in keeping the TM1 and TM2 of the monomer intact on the extracellular side and breaking of this interaction might be crucial for the channel to open, which is also reflected in our analysis. The breaking of D16-Y94 interaction in 5-MTSET probably facilitates the formation of R11-E104 (i,i+1) and E7-R98 (i,i+2) salt bridges, which are discussed in more detail in the following section.

**Fig. 3.**
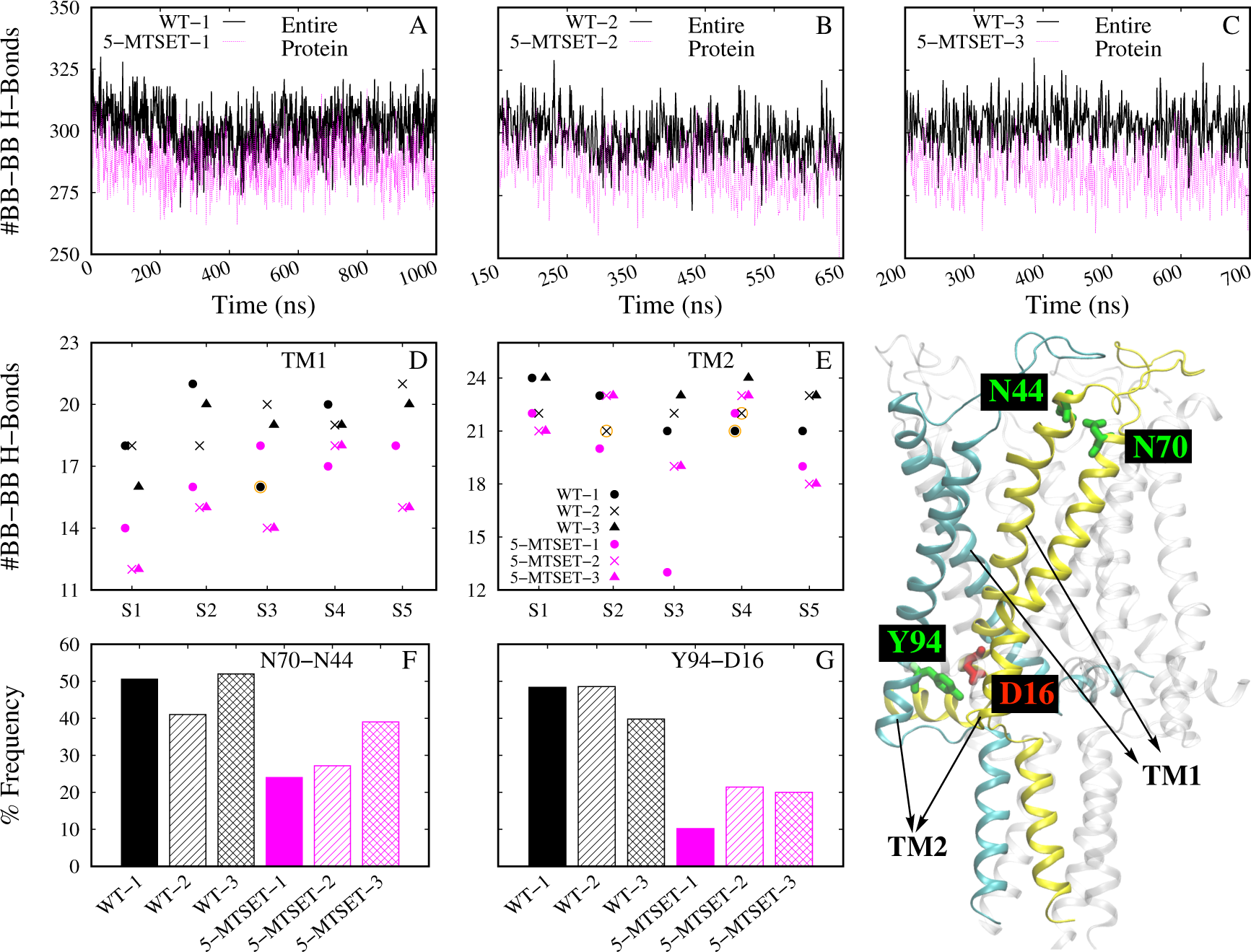
WT vs. 5-MTSET hydrogen bond interaction analysis. (A-C) Comparison of the BB-BB hydrogen bonds of three trials of WT and 5-MTSET respectively. (D-E) BB-BB hydrogen bonds (with > 70% interaction frequency) of TM1 and TM2 of all monomers of WT and 5-MTSET respectively. (F-G) SC-SC interaction frequencies (average of all 5 monomers) of N70-N44 and D16-Y94 in WT and 5-MTSET respectively. D16-Y94 and N44-N70 hydrogen bond interaction pairs are showed in H. D16 (TM1, i) - Y94 (TM2, i+4) interaction pair was on the intracellular side near the labels forming between the TM1 and TM2 of neighboring monomers (i,i+4). N44 (TM1, i)- N70 (TM2, i) was in the extracellular side and forming between the TM1 and TM2 of same monomer (i,i). In the panels A-G all WT and 5-MTSET are represented in black and magenta color respectively. In panels D, E, sets 1,2 and 3 data are represented as circles, crosses and triangles respectively.

### MTSET labels facilitate rearrangement of salt bridge interactions

A salt bridge interaction analysis was done in addition to the H-Bond interaction analysis. Instead of considering the entire trajectory for this analysis, trajectories ranging from 300-500ns and 800-1000ns were examined, respectively (called A and B in Tables 4, 5). Salt bridge interactions are often observed in proteins that give structural and functional conformation and contribute to the recognition, decomposition, catalysis and stability of proteins. ^71^ 2 sets of trajectories were considered in order to understand the stability of these salt bridges. This will provide a good view of salt bridges that were mostly interacting in the simulations. This would provide a clear picture of salt bridge interactions which could be important for protein stability and the conformational difference between WT and 5-MTSET will be clearly classified. In this analysis, 8 interesting salt bridge interactions are identified and categorized as 4 different classes as for the sake of comparison: TM1(i)-TM2(i+1), TM1(i)-TM2(i+2), TM2(i)-TM2(i), and TM1(i)-ECL(i+1). They were also classified into inter and intra-unit salt-bridges. Inter-unit salt bridge interactions are interactions occurring between neighboring monomers/units (TM1(i)-TM2(i+1), TM1(i)-TM2(i+2), and TM1(i)-ECL(i+1)) listed in Table 4). The interaction that are occurring within the same monomer/unit are intra-unit salt-brides (TM2(i)-TM2(i)) listed in the Table 5.

**Table 4:** Inter-unit salt bridge (SB) interactions.

| SB Class | SB | WT |  | 5-MTSET |  |
| --- | --- | --- | --- | --- | --- |
|  |  | A | B | A | B |
| TM1(i)-TM2(i+1) | R11-E104(SB1) | 1/2/1 | 0 | 2/2/2 | 2 |
| TM1(i)-TM2(i+2) | E7-R98(SB2) | 0/0/0 | 0 | 1/1/1 | 1 |
|  | R11-D108(SB3) | 2/2/2 | 2 | 3/1/1 | 1 |
|  | E7-K99(SB4) | 1/2/3 | 2 | 1/1/0 | 1 |
| TM1(i)-ECL(i+1) | K33-D68(SB5) | 1/0/0 | 0 | 1/1/1 | 1 |
|  | D36-R58(SB6) | 0/0/0 | 0 | 0/1/1 | 1 |
|  | R45-D53(SB7) | 2/1/1 | 2 | 0/0/0 | 0 |
| Total |  | 7/7/7 | 6 | 8/7/6 | 7 |

A and B refers to 300-500ns and 800-1000ns trajectories, respectively. Salt bridge interactions with contact frequency $\geq 70\%$ (i.e., the distance between interacting residues was $\leq 4 \text{ \AA}$ for $\geq 70$ of simulation time) were considered. ECL refers to extracellular loops.

**Table 5:** Intra-unit salt bridge interactions.

| SB Class | SB | WT |  | 5-MTSET |  |
| --- | --- | --- | --- | --- | --- |
|  |  | A | B | A | B |
| TM2(i)-TM2(i) | D108-R98(SB8) | 0/1/1 | 2 | 0/0/0 | 1 |
| Total |  | 0/1/1 | 2 | 0/0/0 | 1 |

A and B refers to 300-500ns and 800-1000ns trajectories, respectively. Salt bridge interactions with contact frequency $\geq 70\%$ (i.e., the distance between interacting residues was $\leq 4$ Å for $\geq 70$ of simulation time) were considered.

Out of the 8 significant salt-bridges identified, 7 are inter-unit salt bridge’s and one is a intra-unit salt bridge. The dominating inter-unit interactions in the WT belong to the class TM1(i)-TM2(i+2), followed by TM1(i)-ECL(i+1) and TM1(i)-TM2(i+1). There is no significant difference in the total number of inter-unit salt bridges between A and B section of simulation in the WT (7 vs. 6; Table 4). In the case of 5-MTSET, again the dominating inter-unit salt bridges are TM1(i)-TM2(i+2); followed by the classes TM1(i)-ECL(i+1) and TM1(i)-TM2(i+1), which have equal contribution. Again there is no significant difference between A and B sections of simulations with respect to the total number of salt bridges in 5-MTSET(8/7/6 vs. 7; Table 4). In terms of A and B trajectories salt bridge research, the total number of salt bridges in WT and 5-MTSET are significantly similar. However, there is a difference in distribution of these salt bridges between the WT and 5-MTSET. For example, the number of of salt -bridges belonging to the class TM1(i)-TM2(i+1) in WT is 0, where as in 5-MTSET it is 2 (comparing just B section in either case). There is only one salt bridge belonging to this class (R11-E104; designated it SB1) and it is located on the intracellular side near the bottle neck region (Fig. 4A). Considering that the labels facilitated the SB1 salt-bridge formation in the 5 MTSET (total 2). Since SB1 interactions are so important for protein stability, they play a key role in the activation of the channel pore.

**Fig. 4.**
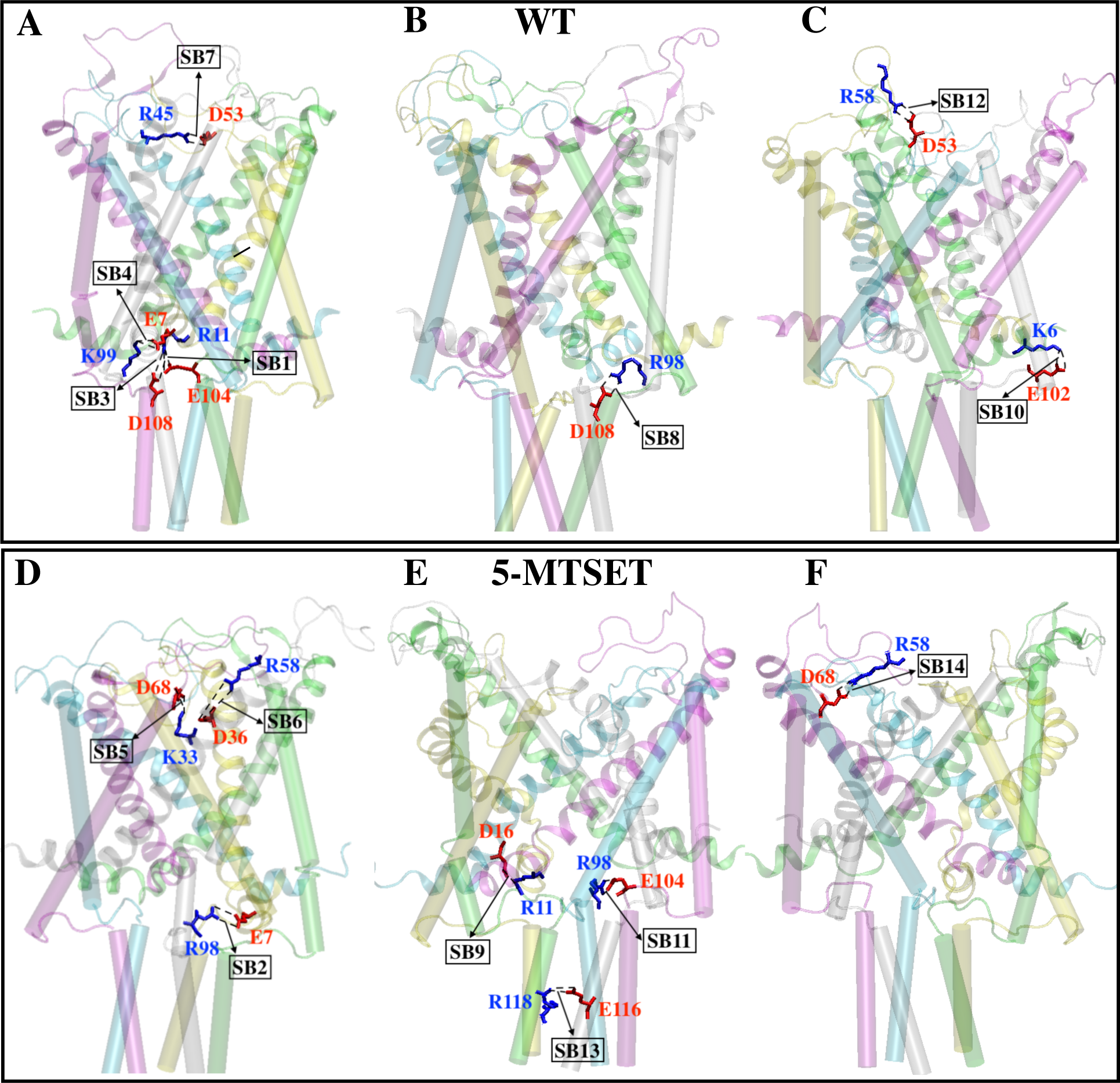
Salt bridge interactions that are playing a key role in functioning of the engineered MscL channel.

On average, there is loss of one salt-bridge belonging to the TM1(i)-TM2(i+2) in 5-MTSET compared to the WT (4 vs. 3; again comparing B in both cases). There is a loss of one SB3 (belongs to TM1(i)-TM2(i+2) helix between R11-D108), one SB4 (belongs to TM1(i)-TM2(i+2) helix between E7-K99), and gain of one SB2 (belongs to TM1(i)-TM2(i+2) helix between E7-R98) in the 5-MTSET. Out of these three mentioned above, SB2 does not exist in the WT at all. SB2 could be a possible salt bridge that was facilitated by labels in opening of channel. With respect to the TM1(i)-ECL(i+1) class, both the WT and 5-MTSET have two salt-bridges each. However, the distribution is different in both cases; WT has only SB7 (R45-D53) belonging to this class, which is completely absent in the 5-MTSET (2 vs. 0). There is a gain of one SB5 and one SB6 in the case of 5-MTSET, which are completely absent in the WT. Therefore, the difference in the distribution of TM1(i)-ECL(i+1) salt-bridges between WT and 5-MTSET is crucial for the complete opening of the MscL channel pore. Particularly, the SB7 interaction ties the neighboring monomers/units on the extracellular side. Hence, the loss of this interaction on the extracellular side is key for the opening of the channel. It is this SB7 interaction that keeps the loops in the outer circumference. In the WT, there are 2 of these SB7 salt bridges keeping the channel intact, where as these interactions are lost (total 0) in the 5-MTSET allowing the loops to freely move and facilitating their collapse into the center of the protein. On the other hand, the SB5 and SB6 interactions that are formed in the 5-MTSET (completely absent in WT) on the extracellular side facilitates the loops collapsing into the center of the protein.

Based on above factors one can conclude that the sudden jerk-like motion induced by the labels at the bottleneck region facilitates the complete loss of SB7 in 5-MTSET, which in turn facilitates the formation of salt-bridges SB5 and SB6 (Fig. 4D). We believe that the salt-bridges SB5 and SB6 are blocking the complete opening of the engineered MscL channel on the extracellular side. Hence, the loss of these 2 salt-bridges (SB5 and SB6) is very important, which did not happen in our unbiased MD simulations within the accessible timescale. Based on these finding, one could believe that the gain of two TM1(i)-TM2(i+1) and loss of one (TM1(i)-TM2(i+2) salt-bridge interactions facilitated the opening of the channel on the intracellular side; and gain of SB5 and SB6 on the extracellular side prevented the opening on the extracellular side.

Only one intra-unit salt-bridge was identified in our simulations belonging to the TM2(i)-TM2(i) helix formed between D108-R98 (denoted as SB8). There is a loss of this intra-unit salt-bridge interaction (SB8) (Fig. 4B) belonging to the class TM2(i)-TM2(i) in 5-MTSET (2 vs. 1; Table 5). Probably the loss of this particular interaction in 5-MTSET, which is at the intersection of TM helices and the intracellular helices (IH), might have made the TM helices and the corresponding IHs more flexible allowing the opening of the channel.

### Interhelical angle analysis confirms the partial spontaneous opening of engineered MscL

To understand the mechanism of opening of the channel in response to the introduction of positively charged MTSET labels, various inter-helical angles were measured, which are defined as alpha (α), beta *((3),* gamma (r) and delta (b) as discussed above (Figs. 1 B and D). The overall behavior of the inter helical angles (Fig. 5) of all 5 subunits (S1, S2, S3, S4, and S5) in WT are very uniform, in each case in WT are very uniform, except in a very few limited cases, for example S4-S5 *γ* orientation angle (Fig. 5C). However, in 5-MTSET simulations the orientation angles show very distinct behaviour when compared to WT. All five inter-helical angles are very dispersed and more deviating from the initial structure in every orientation angle (*α*, *β*, *γ*, and *δ*) compared to WT (Fig. 5A-D). Although there is a significant difference in few cases between the WT and 5-MTSET, such as S2-S3 *α* (Fig. 5A), S2-S3 *β* (Fig. 5B), S2-S3 *γ* (Fig. 5C), and S1 *δ* (Fig. 5D), overall a uniform behavior with respect to 5-MTSET in any of the 4 inter-helical angles are not observed in these findings. This behavior probably is due to the fact that 5-MTSET is not completely open in the equilibrium simulations.

**Fig. 5.**
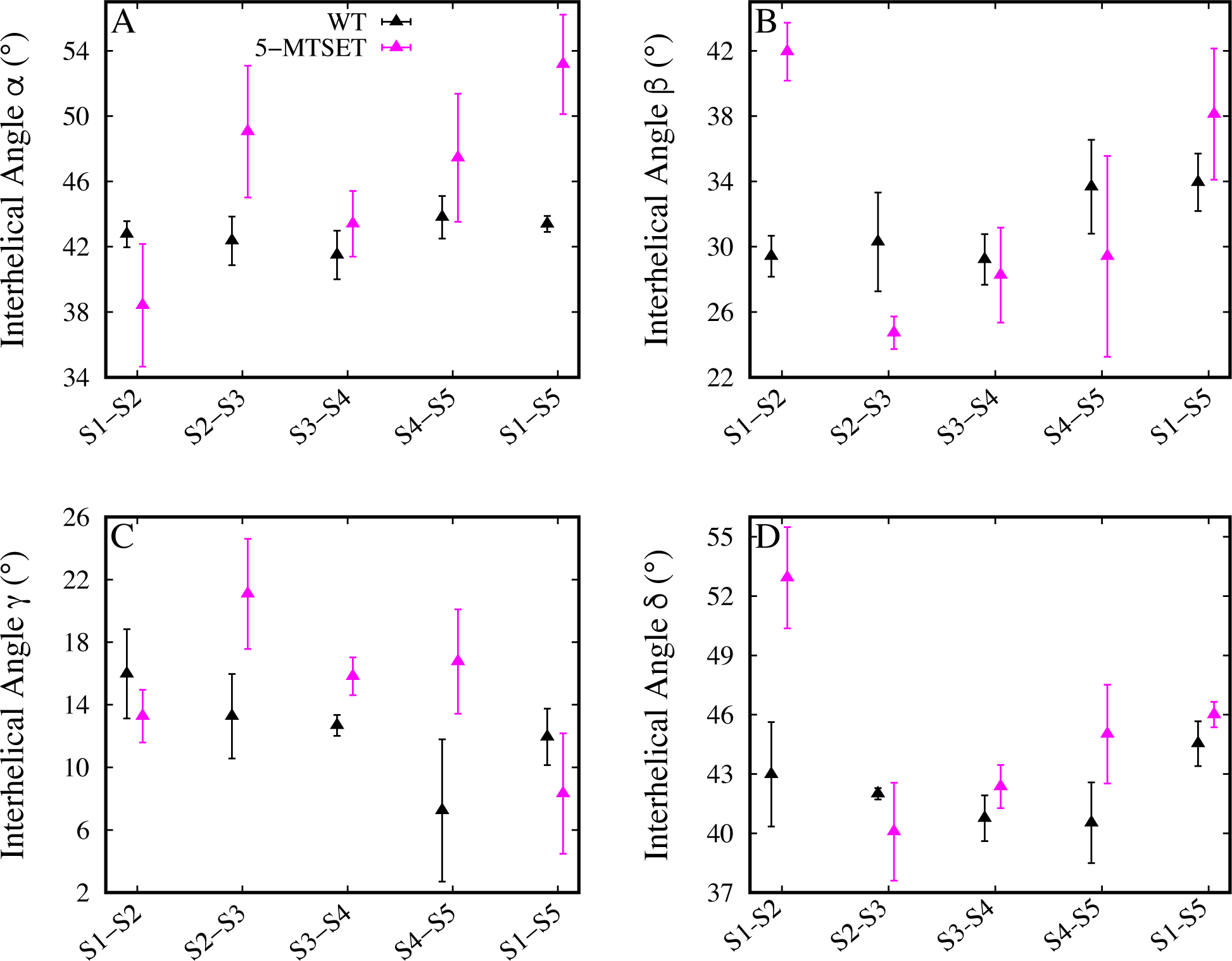
Orientation angles *a, {3,* /,and *b.* (A-D) Comparing the inter-helical angles *a* (A), *f3* (B), 1 (C) and *b* (D) between the WT and the 5-MTSET respectively. WT and 5-MTSET data points are represented as black and magenta triangles res pectively. Each data point represents the average and standard deviation of all three simulation sets in each case. Only last 300ns of trajectory data is considered for this analysis.

### Principle Component Analysis confirms that loops play a key role in the dynamics of MscL channel

To unveil the principle variatio ns between the WT and engineered structures, principle component analysis (PCA) was performed. Usually the top ranking modes reflect the dominant features responsible for the variations in the structures. Projections onto principle components PC1 and PC2 clearly discriminate WT, 1-MTSET and 5-MTSET systems (Fig. 6A). The contribution of these two PCs to the total variance is 51.4%. Top 5 PCs’ contribution to the variance is 64.3% and the top 15 PCs’ is 81.4% (Fig. 6D). Although, the deviation of the engineered systems from the WT is not huge, all three systems are clustered differently along PC1 space. The 5-MTSET systems were found to be deviated more than the 1-MTSET systems from the WT.

**Fig. 6.**
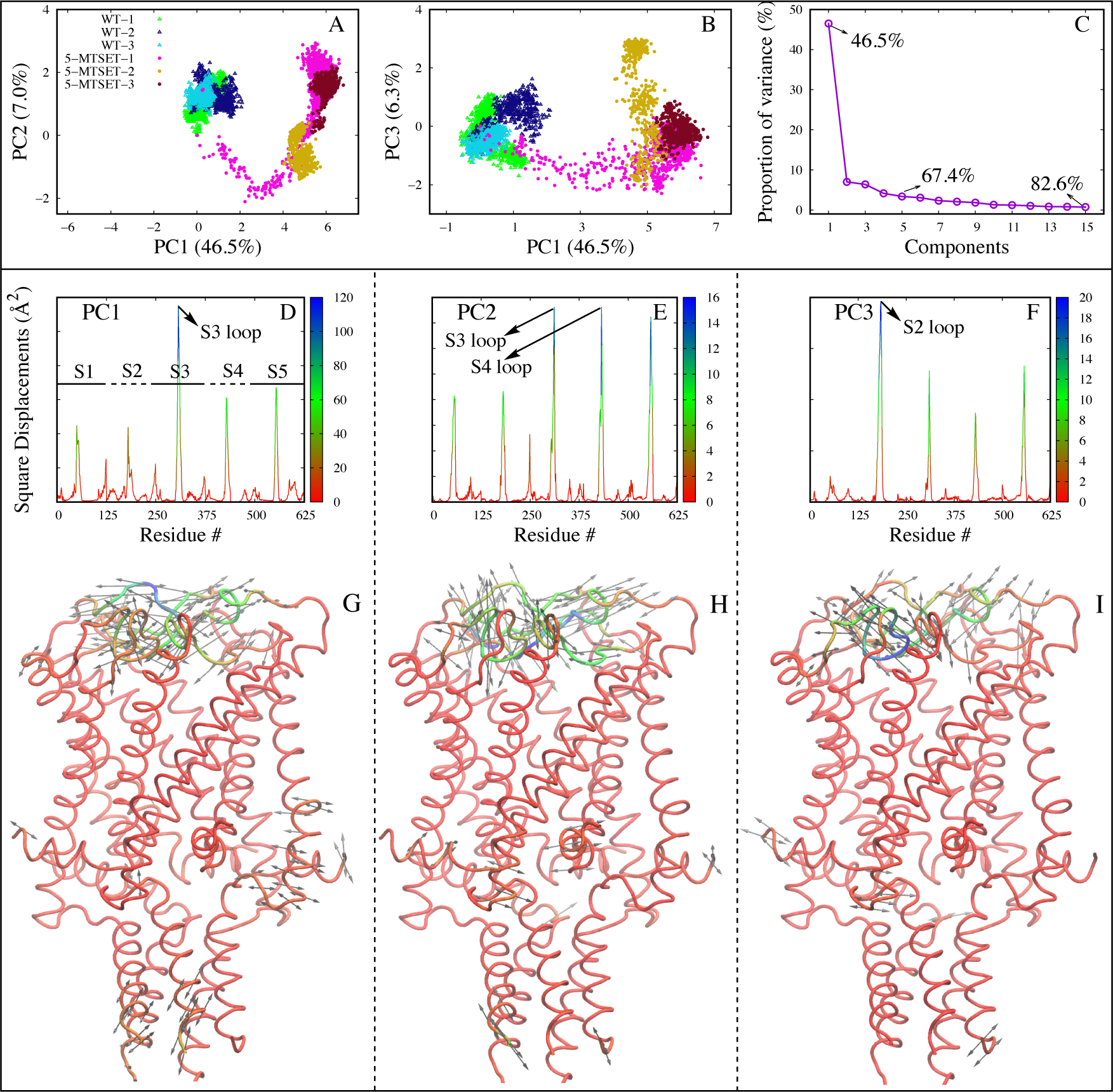
Pro jections alo ng PCs 1, 2 and 3. (A) PC1 vs. PC2. (B) PC1 vs. PC3. (C) is the pro portion of variance (or total mean square displacement) of first 15 PCs. (D-F) Co ntribution of individual residues (represented as mean square fluctuations of the atom positions) for PCs 1, 2 and 3 respectively. (G-I) Structural variations along PCs 1, 2 and 3 res pectively. Structures are colored based on the extent of mobility/fluctuation. Bidirec­ tional arrow s hows the direction of the fluctuation of the structure and the length of the arrow reflects the magnitude of fluctuation.

All the triplicate simulations of each system were clustered very close to each other, elucidating the fact that the structural variations along PCs 1 and 2 are reproducible and significant. PC2 separates 1-MTSET from 5-MTSET and WT systems (Fig. 6A); PC3 and PC4 show how dispersed the 5-MTSET system is compared to the WT and 1-MTSET systems (Figs. 6 B and C). The structural variation along PC1 is dominated by the loops in S3 monomer (blue colored peak in Fig. 6A) followed by the loops in S4 and S5 monomers (orange and yellow peaks in Fig. 6A). Overall, key outcome of PCA analysis is that the loop behavior of WT and engineered MscL protein are significantly different. For example, the residue with the highest fluctuation in PC1 is R58 belonging to loop S3 (*β* = 78.6Å). R58 is involved in salt bridge interactions with D36 and D68 as demonstrated earlier. And the PCA analysis supports the earlier salt bridge interaction analysis and the hypothesis that the loops are playing a key role in the dynamics and function of MscL channel. The second most dominant source of fluctuations after the loops are the intracellular helices, especially S1 and S2. In the PC2, S1 loop dominates (purple peak in Fig. 6B, residues 50-56) followed by S5 loop (yellow peak in Fig. 6B, residues 56-60). In PC3, again loops dominate and the contribution of all loops is approximately similar (Fig. 6C) followed by loops in the intracellular helix of the S2.

### Obtaining the open MscL structure via non-equilibrium simulations

Since the unbiased equilibration simulations failed to fully open either the WT or the engineered MscL structures as explained earlier, non-equilibrium (NE) pulling simulations simulations are required to induce the full opening of the resulting unbiased equilibrium simulation structures. To open these systems, a previously created model of WT active /open MscL based on smFRET data was used as a reference. ^68^ Orientation collective variables (colvars) were used to steer the initial structures towards the target. ^72, 73^ Where a time-dependent biasing potential is used to move the protein through a distance or rmsd based reaction coordinate from one state to another. The key difference of the methodology discussed here is that a large fraction of analytical efforts are concentrated on discovering appropriate system-specific reaction coordinates and functional biasing protocols to build more efficient transfer pathways In particular, several collective variables were modeled based on ”orientation quaternion” ^50, 67^ formalism as an efficient method to induce rotational transformations in a molecular structure. The steered molecular dynamic (SMD) procedure for WT, 5-MTSET and 5-MTSET(*) structures are done to drive the system from known closed conformation state to open state ^68^ using orientation collective variables as explained above. The SMD simulation was 100 ns long and in each case the SMD procedure was repeated multiple times to support our results.

The work required to open these different structures in NE simulations was compared and assessed to verify which system needs higher work to be activated/opened using the SMD technique. As expected WT structure requires more work to open compared to the engineered 5-MTSET structure (Fig. 7A) and this same behavior was observed in our SMD repeats which supports our hypothesis. To check whether the stating orientation of labels has an impact on the work required for opening the channel, the SMD simulation was done with 5-MTSET(*) structure. The work profile values between 5-MTSET and 5-MTSET(*) structures are significantly similarly (Fig. 7A), leading us to conclude that there is no impact of the starting orientations of the labels on the opening/activation of the channels. The NE work trends support earlier observations that labels facilitate the opening of the channel. Since these structures were force opened, irrespective of the presence or absence of labels all the structures were opened to the same extent (Fig. 7C), which is reflected in the RMSD calculations (Fig. 7B). Although there is a difference in RMSD between the WT(1) and the 5-MTSET(*) set-1 simulations at the start, by the end of 100ns NE simulation there is no significant difference. However, the extent of opening of the pore near the bottle neck region (z= -5 to -10; the structure sort of flattened while opening, hence the bottle neck region went slightly up compared to the equilibrium simulations) is greater in WT(1) compared to the 5-MTSET(*). This is evident from the water content calculations, where WT(1) has more water in the pore region than the 5-MTSET(*)(7 vs. 22 near the narrowest part of the pore, which is Z = -7 Å) (Fig. 7E). This can be because of the labels which are predominantly oriented parallel to the membrane in the earlier equilibration simulations, and so serve to obstruct the water entry (Fig. S1B). However, in the NE simulations, they are most likely oriented perpendicular to the membrane normal (Fig. 7G), and thus obstruct the pore.

**Fig. 7.**
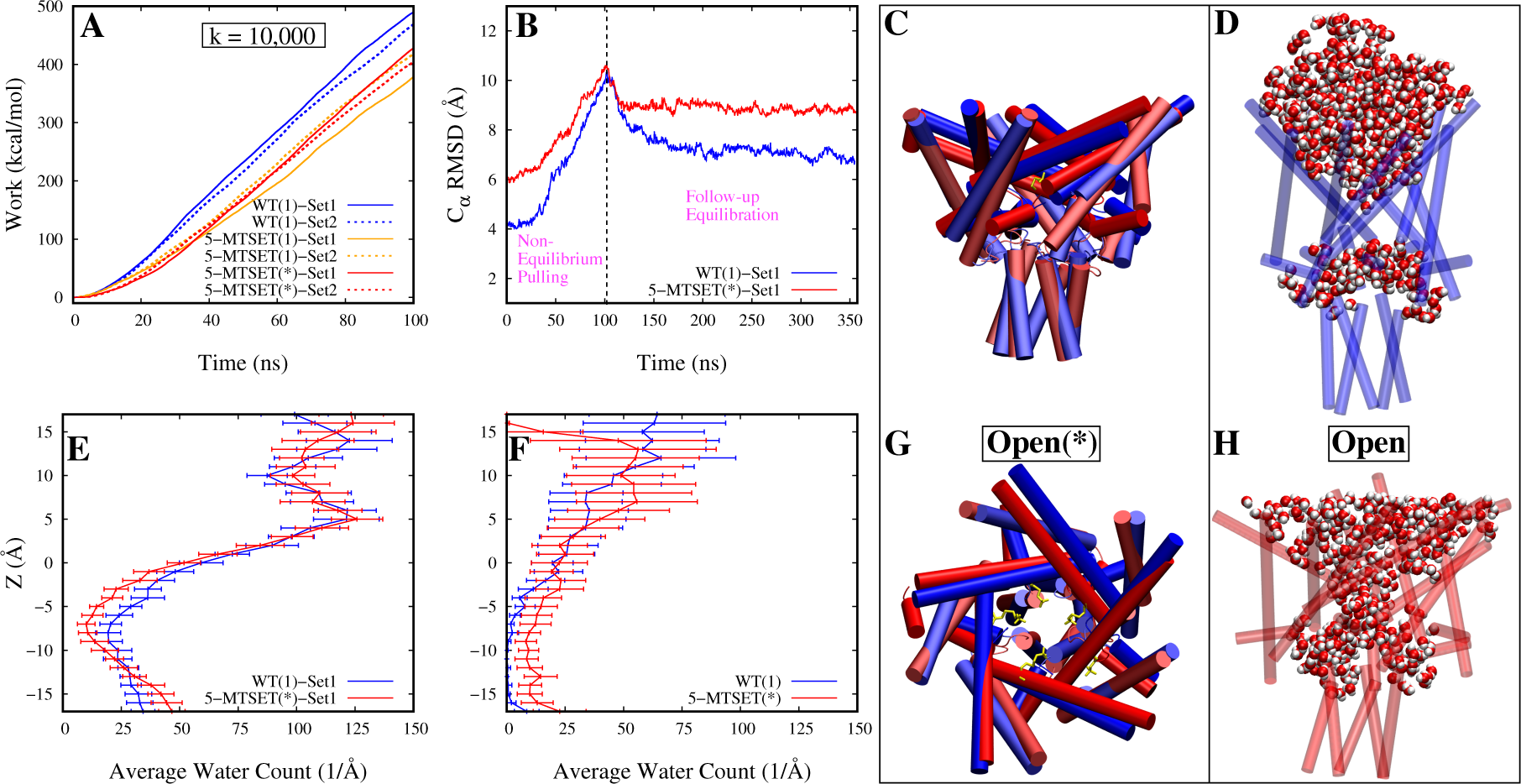
Non-equilibrium and follow-up equilibrium simulations res ulted in an open MscL structure. (A) Comparison of WT vs. engineered MscL (5-MTSET and 5-MTSET(*)) work values as a function of time. (B) Protein Co: RMSD vs. time during the lOOns non­ equilibrium pulling simulation and the follow-up 260ns equilibrium relaxation simulation of WT(l) and 5-MTSET(*) systems. For calculating the Co: RMSD of the protein, crystal structure was used as reference. (E-F) Water count across the pore resulting from lOOns non-equilibrium simulations for WT(l) and 5-MTSET(*) systems, and F is water across the pore during the 260ns follow-up equilibrium simulations. Last 2.5ns simulation was used for the water content calculation. (C) Superposition of open(*) structures of WT(l) and 5-MTSET(*) resulting from non-equilibrium simulations. WT(l) and 5-MTSET(*) are represented in blue and red colors res pectively. (G) Same as *C* except that *C* is side view and G is the top view. (D,H) Open structures of WT(l) and 5-MTSET(*) structures resulting from follow-up equilibrium simulations. Water within the 3Aof the protein is s hown both cases.

### Characterizing the activation mechanism of engineered MscL using SMwST

SMwST simulations were used to further characterize the NE simulation findings in order to examine the conformational dynamics and convergence of the minimum free energy pathway of channel opening. This method is carried out in iterative set of small simulations. This study reports the application of SMwST on conformational changes in the bio-molecules based on atomic level simulations. Starting with an approximate channel opening conformational changes captured in an order of close state (CS) to open state (OS) using a set of 50 images from NE simulation. Optimization of CS to OS was carried out by applying a set collective variables (10 quaternion colvars). Different variable PCA analysis was done to reveal the principle modifications over the course of each iteration (180 iterations). The superior features accounting for the differentiation in the structure are reported in the top ranking modes. The results of the PC1 and PC2 clearly shows difference in optimization process from CS to OS (Fig. 8C) for every image. These two PC’s contribute a total variance of 72.3%. The reported structural differentiation over the PC1 is the result from the activation/opening of the MscL at different states and PC2 is the result from the optimization for each iteration. All three PC’s contribute a variance of 76.0%(Figs. 8 C and D). A PCA analysis was also implemented on our quaternion(QPC) variable used for convergence of CS to OS in SMwST for every iteration. These reported QPC’s 1 and 2 contribute a total variance of 96.73% (Fig. 8B). Projected QPC’s are reported for every iteration show converging of the opening mechanism. The dissimilarity of the QPC1 values are resulted from iterative SMwST simulations aimed at improving the sampling by using progressively more optimal reaction coordinates and more relaxed conformations. This shift from right to left (Fig. 8B) on QPC1 is due to the relaxation of the structure in each iteration which captures a smooth and efficient pathway of the MscL activation. The superimposition of last frame of iteration 1 and 180 (blue and red respectively (Fig. 8A)) shows how divergent both structure are, where MscL at iteration 180 (Fig. 8A(red))has more open and relaxed conformation compared to the iteration 1 (Fig. 8A(blue)). However our observations of SMwST optimization of channel opening have not converged completely even after an extensively long simulation.

**Fig. 8.**
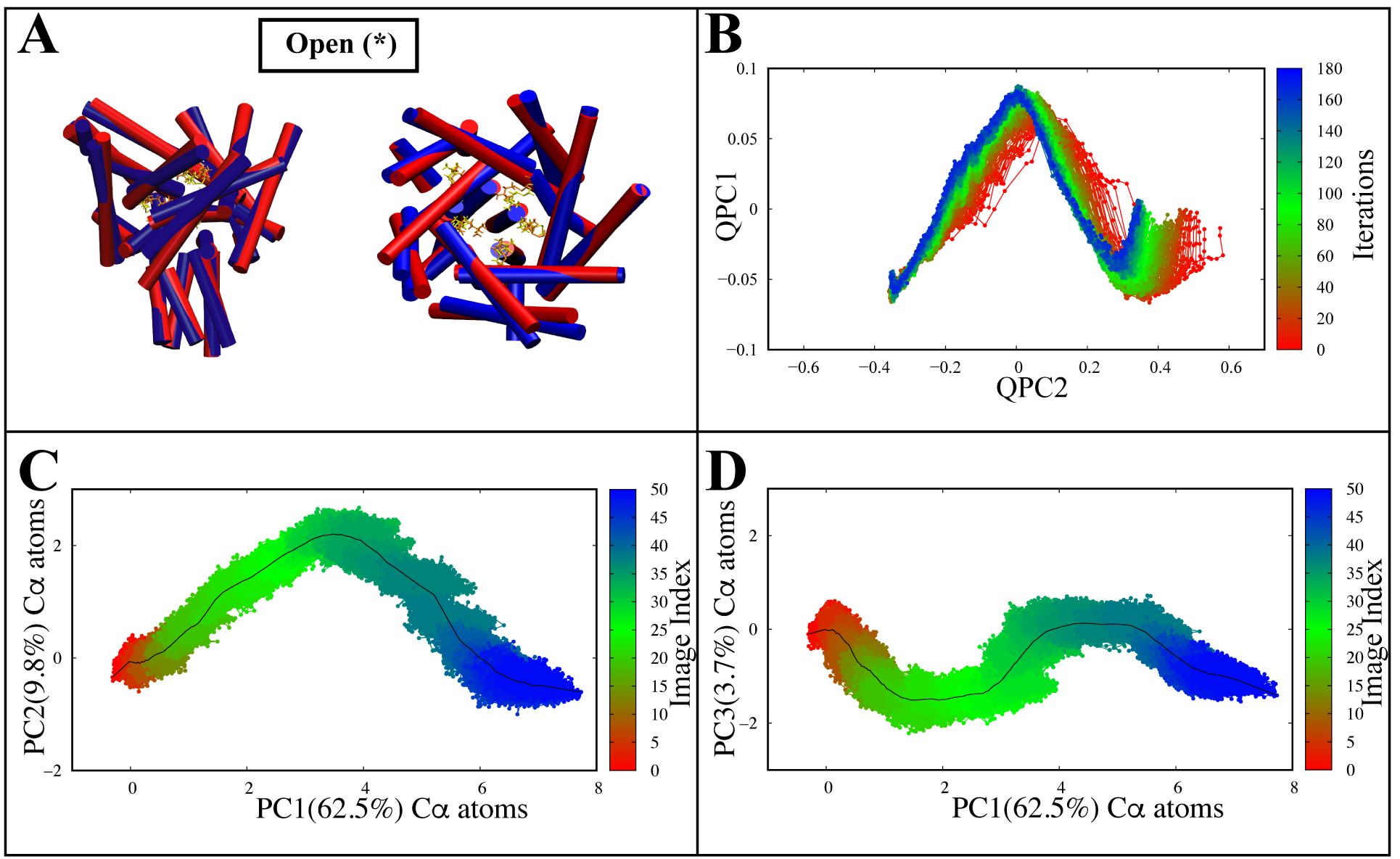
Characterizing the activation mechanism of engineered MscL using SMwST. (A-D) superposition of iteration 1(blue) and iteration 180(red) resulting from SMwST(A), QPC1 vs. QPC2 that is, the principal components of the vector parts of the Q quaternions of SMwST (B), Comparison PC1 vs. PC2 of alpha carbons (C), Comparison PC1 vs. PC3 of alpha carbons (D), All 180 iterations of SMwST were considered for this calculation.

**Fig. 9.**
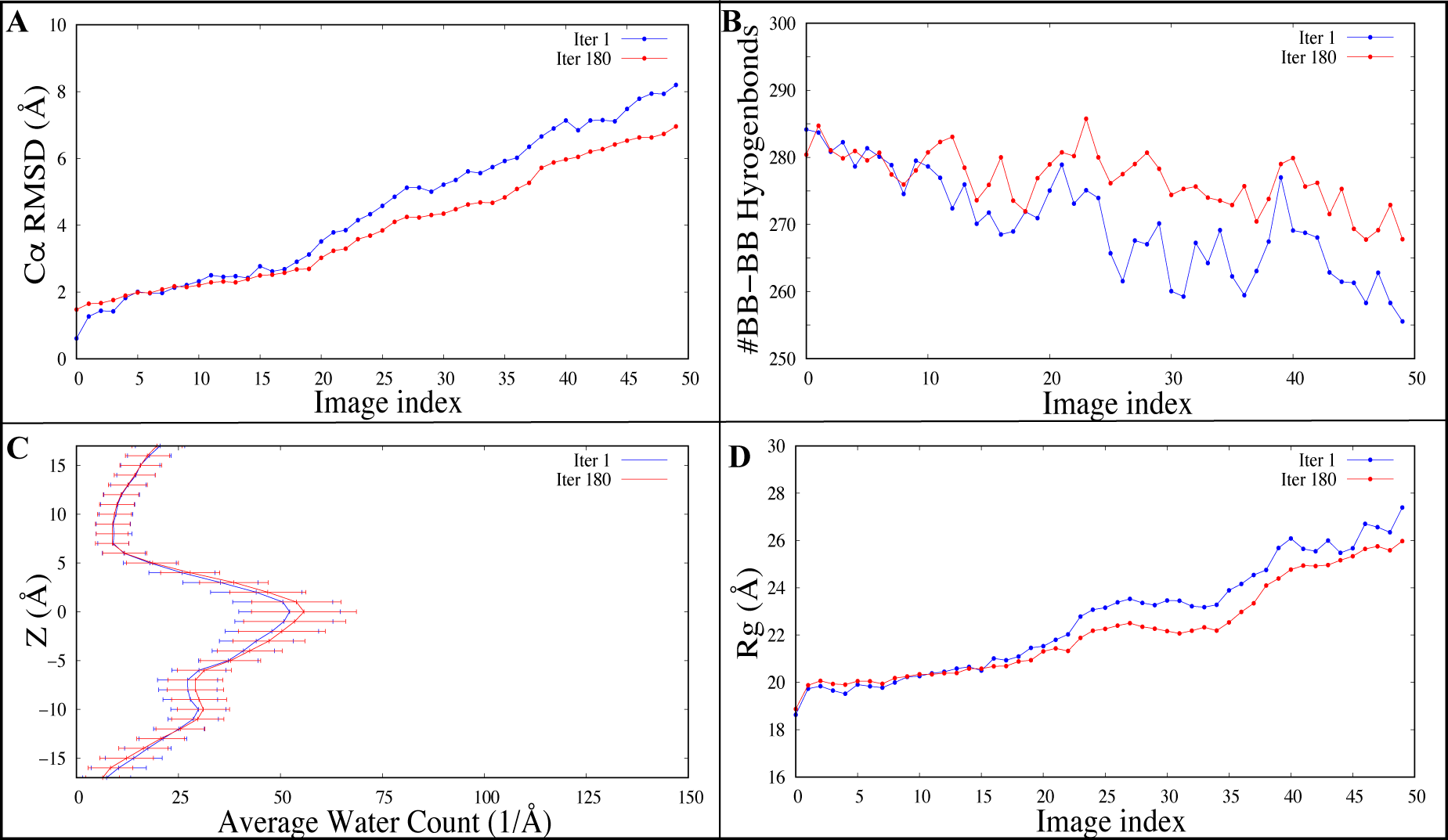
String method simulation analysis of RMSD, number BB-BB hydrogen bonds, average water count and radius of gyration(Rg)· (A-D) protein Co: RMSD (A), number BB-BB H-bonds (B), average water count across the pore (C) and radius of gyration *(Rg)* between iterations 1 vs. iterations 180 systems. Iterations 1 and Iterations 180 data are represented in blue and red respectively. The complete data of iteration 1 and 180 are considered for the above calculation and each point value is averaged over 20 copies of each image.

The following variables were analyzed to validate our divergent PC analysis and explain the variations between iterations: BB-BB H-bonds, RMSD of alpha carbons, average water count and radius of gyration *(Rg)* on first and last iteration trajectories of SMwST for all 50 images each. Each images consist of 20 parallel set of SMwST simulations as discussed above. Based on the previous results of targeted MD simulation, the breaking of hydrogen bonds play a major role in activation of the protein. Analysis H-bonds in process of channel opening would be a best choice to show iterative optimization between iteration 1 and iteration 180, in which the dissociation of H-bonds in iteration 180 occurs in a distributed and relaxed manner whereas iteration 1 reports noisy and rapid separation of bonds. Same results are replicated in RMSD of alpha carbons, average water count and radius of gyration where MscL in iteration 180 is activated with more relaxed conformations than in iteration 1 (Figs. 9 A, C, and D). This above extensive analysis provide an explanation for the iterative relaxation of SMwST method. All above results correlate with each other.

Since SMwST simulations have not converged comprehensively as expected we further analysed our NE structures. To check the stability of the non-equilibrium structures, the set-1 of WT and 5-MTSET(*) non-equilibrium structures were relaxed via the unbiased equilibration simulations using the protocol explained in Table 2. In these follow-up equilibration simulations (FE) both the WT and 5-MTSET(*) structures were not stable, WT collapsed back to the original state while the 5-MTSET(*) collapsed to a state which is more open than the equilibrated structures (we call it open) (Fig. 7 D and H). This also lead us to hypothesize that the open/active state of MscL might not be as open as previously believed. This is reflected in the water content calculations, where the average number of waters in the pore near the bottle neck region is 0 in the WT, and ≈ 10 in 5-MTSET(*) (Fig. 7F).

The water content correlate well with the C*α* RMSD (Fig. 7B).

### Characterizing the open MscL structure resulting from the followup NE simulations

The WT(1) and 5-MTSET(*) MscL structures resulting from the follow-up NE simulations were further characterized. To compare and contrast the WT(1) and 5-MTSET(*) open structures, variables like RMSF, ECL Rg, number of BB-BB H-Bonds, average Y94-D16 H-Bond frequency, SC-SC (both inter-/intra-unit) salt bridge interactions, and different interhelical angles were analyzed. Significant differences between the open WT(1) and open 5MTSET(*) structures were observed. The overall fluctuation of the 5-MTSET(*) structure is greater than the WT(1) (Fig. 10A). The *R_g_* of ECL of both the structures was reduced for the entire length of the trajectory. However, the *R_g_* of 5-MTSET(*) is significantly greater than the WT(1) (Fig. 10b); there is at least 4 Å difference between the two at the end of the 260ns follow-up NE simulation, which shows that the ECL region of 5-MTSET(*) is more open compared to the WT(1). This is in contrast to the equilibration simulations, where the *R_g_* of the 5-MTSET structures are lower than the WT (Figs. S3 D and F). One can conclude that this is due to the formation of SBs 5 and 6 in the equilibrated 5-MTSET structures (Table 4) as explained earlier, and that these SBs were not formed in the 5-MTSET(*) structure in the follow-up NE simulations (Table 6). Similar to the equilibrium simulations, the number of BB-BB H-Bonds in 5-MTSET(*) is greater (on average 15) than in the WT(1); although they were similar at the start of the simulations, as the simulations proceeded the difference became significant (Fig. 10C). This establishes the fact that the loss of BB-BB H-Bonds makes the structure more flexible in this region by facilitating the opening/activation. However, this trend is not clearly visible in the TM region. For example, the number of TM1 BB-BB H-Bonds of S1, S2, and S3 in WT(1) are greater than 5-MTSET(*), where the opposite is true for S4 and S5 (Fig. S4E). On the other hand, the number of TM2 BB-BB H-Bonds of S1 and S2 in 5-MTSET(*) are greater than WT(1), where it is opposite for S3, S4 and S5 (Fig. S4F). Also, there is a loss of SC-SC Y94-D16 H-Bond in 5-MTSET(*) (*≈* 15 frequency%) compared to the WT(1) (*≈* 45% frequency); this behavior is similar to the equilibrium simulations (Fig. 10D). This supports the hypothesis that the loss of this particular H-Bond is key for the activation of engineered MscL channel.

**Fig. 10.**
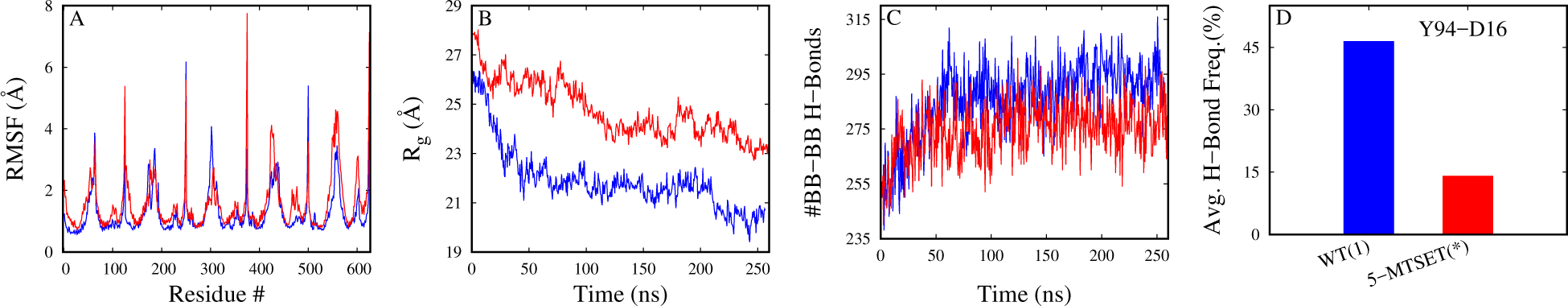
Characterizing the open MscL structure. (A-D) Comparison of RMSF (A), radius of gyration (B), number of BB-BB H-Bonds (C), and Avg. Y94-D16 H-Bond frequency (D) between WT(1) vs. 5MTSET(*) systems. WT(1) and 5MTSET(*) data are represented in blue and red, respectively. The entire 260 ns of follow-up NE simulations data is considered for this calculation.

**Table 6:** Inter-unit salt bridge interactions.

| SB Class | SB | WT(1) | 5-MTSET(*) |
| --- | --- | --- | --- |
| TM1(i)-TM2(i+2) | R11-D108(SB3) | 2 | 1 |
|  | E7-K99(SB4) | 1 | 0 |
| TM1(i)-ECL(i+1) | R45-D53(SB7) | 1 | 0 |
| TM1(i)-TM1(i+1) | R11-D16(SB9) | 2 | 1 |
| TM1(i)-TM2(i+1) | K6-E102(SB10) | 3 | 0 |
| TM2(i)-TM2(i+1) | R98-E104(SB11) | 2 | 1 |
| ECL(i)-ECL(i+1) | D53-R58(SB12) | 1 | 0 |
| IH(i)-IH(i+1) | E116-R118(SB13) | 2 | 1 |
| **Total** |  | 14 | 4 |

Only last 100 ns of relaxation simulation trajectory is considered. IH refers to intracellular helices. Interactions with >70% are considered. SB is salt bridge.

Apart from this, the SC-SC SB interactions (both inter-/intra-unit) were studied. Unlike the earlier equilibrium simulations, the total number of inter-unit SBs in the case of 5-MTSET(*) is 4 (i.e., the SBs with ≥ 70% interaction frequency), where as in the case of WT(1) it is 14. Overall, 5-MTSET(*) has to loose 10 inter-unit SBs compared to the WT)(1) for it to open in the follow-up NE simulations. These 14 inter-unit SBs identified in the WT(1) is classified into 7 different classes as shown in Table 6. Of the 7, two were already introduced earlier (TM1(i)-TM2(i+2) and TM1(i)-ECL(i+1); Table 4) and the remaining 5 are introduced here since these classes of SBs were not found earlier in the equilibrium simulations in either WT or 5-MTSET structures. The 14 WT(1) SBs are spread all over the channel, however, the dominating interactions are between the TM1 and TM2 of the neighbor units (total 6 of 14). It is very interesting that 14 were found in WT(1) in the follow-up NE simulations, where as only 6 are identified in the earlier equilibration simulations. This is probably because the gain of 8 new SBs in the WT(1) is needed for the WT(1) protein to collapse back to the original state, whereas in the case of 5-MTSET(*) there is a overall loss of ≈ 3 SBs compared to 5-MTSET in the earlier equilibration simulations (≈4 vs. ≈7).

The two classes of SBs that were observed in the 5-MTSET in the equilibrium simulations, i.e. TM1(i)-TM2(i+1) and TM1(i)-ECL(i+1), are completely lost in the NE follow-up equilibrium simulations. This supports our earlier hypothesis that the loss of TM1(i)-ECL(i+1) SBs are key for the complete opening of the engineered channel. There is not a single SB involving ECL in the case of 5-MTSET(*), which further supports our hypothesis. However, there is gain of 3 new classes of SBs in 5-MTSET(*) in NE follow-up equilibrium simulations: TM1(i)-TM1(i+1), TM2(i)-TM2(i+1), and IH(i)-IH(i+1). Additionally, there is a gain of 2 new intra-unit SBs in 5-MTSET(*) (Table 7) and which belong to the class ECL(i)-ECL(i) (another new class that was not found in the earlier equilibrium simulations). These two SBs are also not observed in the WT(1).

**Table 7:** Intra-unit salt bridge interactions.

| SB Class | SB | WT(1) | 5-MTSET(*) |
| --- | --- | --- | --- |
| ECL(i)-ECL(i) | D68-R58(SB14) | 0 | 2 |
| **Total** |  | 0 | 2 |

Only last 100 ns of relaxation simulation trajectory is considered.

To illustrate the partial closing of the protein (Fig. 11) that was observed in follow-up NE simulation. Principle components analysis was done on FE simulations plotted against the SMwST structure (Fig. 11). Projection of QPCs on unbiased simulations and SMwST results (Fig. 11B) represent the global conformational transformation of protein from OS to partial OS state in 260ns simulation period. One might expect this to be an intermediate transition state since our observations shows that the protein was very stable at this state for a long period of our simulation. While all our global conformational transition PCA results are in agreement with each other (Figs. 11 C and D). This dissimilarity over QPC1 (Fig. 11B) and PC1 (Figs. 11 C and D) from right to left are the results of the conformational transition from OS to intermediate OS conformation as discussed above.

**Fig. 11.**
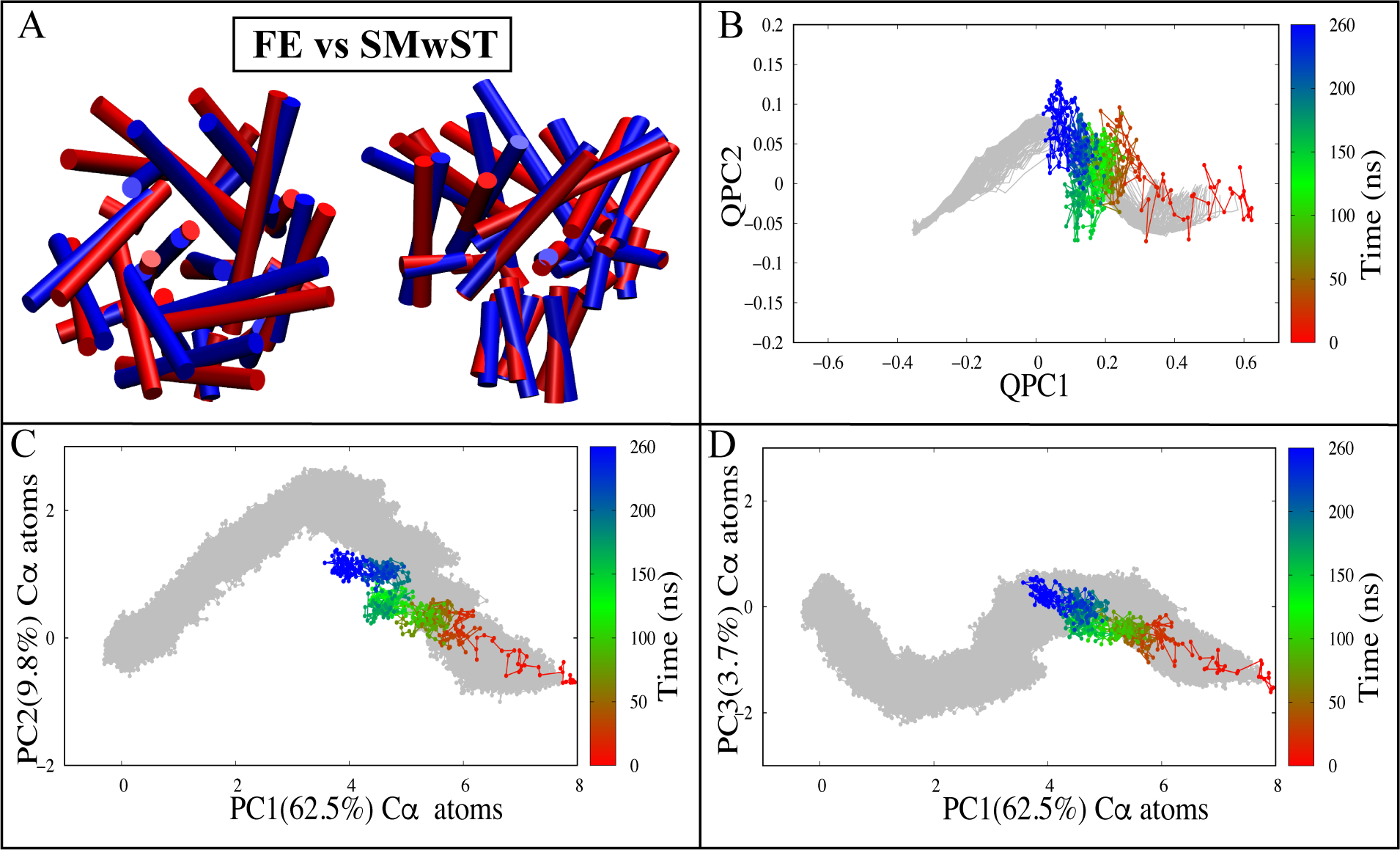
Follow up equilibrium simulation analysis (A-D). (A) Superposition of last frame of FE(blue) and SMwST (red), (B) QPC1 vs. QPC2 of FE simulation, (C) PC1 vs. PC2 of alpha carbons, (D) PC1 vs. PC3 of alpha carbons. The entire 260 ns trajectory of FE simulation data is considered for this calculation. SMwST simulation results (Fig. 8) are projected in the background (gray).

Each image/window was equilibrated post SMwST method, to obtain free energy landscape of each state associated with activation of the MscL. These calculations were carried out using Expectation Maximization (EM) algorithm for different sets of generator matrices. The generator matrix is constructed from, a number of observations of MscL state, where state i is considered to be the state at time t, state j is considered to be the state at time t + *T,* and *T* is a given lag time. The SMsST was used to sample the phase space and discover important reaction coordinates to generate the matrix. From this transition matrix the rate/generator matrix was created using EM algorithm and substitute the values in free energy calculation. The methodology described in Goolsbyet. *al.,*^70^ is used to quantify EM and free energy. The calculation of average free energies are done using first 10 *τ* reported at low lag time. Figure 12 shows the estimated free energy calculation of each conformation at various degrees of openness. These results suggest that during CS -----+ OS transition, the free energy of initial open state and final closed conformation state are higher than any intermediate conformation allowing protein inter-conversion between these two states. The MscL protein activation free energy landscape begins with a high-energy CS state followed by a steady decline to a lower free energy sate and then a climb back to a higher-energy open state. All together, the free energy of CS and OS are much higher than any transition states along the activation pathway. One might postulate these two states as being very unstable, leading to the collapse of OS protein back to a transition intermediate state. This is exactly what we observe in our FE simulation results (Fig. 11A-D), where the final structure from NE simulation (OS) was equilibrated for 260ns resulting in a partially open state. The findings of the free energy landscape are in agreement with our previous postulates concerning the, conformational transition from OS to intermediate OS state observed in FE simulation.

**Fig. 12.**
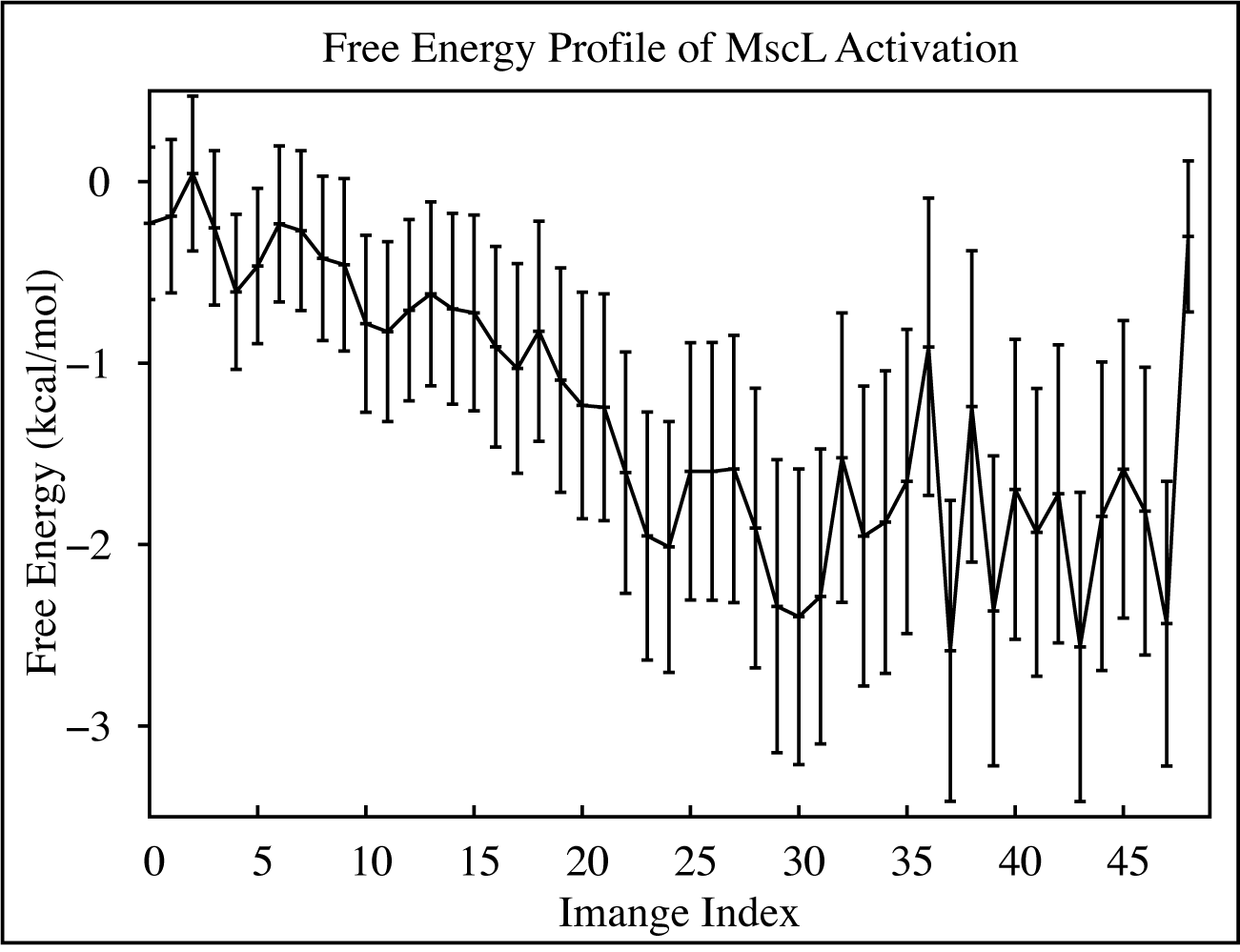
Free energy profile of each conformation in MscL activation. The figure represents the average of first 10 *T* free energy of MscL activation for each image/window. Each image represents a particular conformation of MscL in activation pathway composed of 50 images. Error bars represent standard deviation of free energy calculated for first 10 *T* generator matrix.

### Proposed mechanism of action

Based on our detailed analysis of the equilibrium, non-equilibrium, swarm-based string method and follow-up equilibration simulation data, a hypothesis for the opening/activation of the engineered MscL channel is proposed in this extensive study. According to the hypothesis, first the channel opens spontaneously due to the repulsion between engineered positively charged labels and also due to steric clashes between the labels and other bottle neck residues, which is also demonstrated through interaction analysis (8 among the top 10 residues interacting with the labels are non-polar (Table S1)). This sudden jerk like motion near the bottle neck region leads to the breaking of inter-unit SB7 interaction (TM1(i)-ECL(i+1)) on the extracellular side, freeing the loops and allowing them to collapse to the center of the protein (depicted by the reduction of radius of gyration (*R_g_*)) (Figs. S3 C and D), which in-turn is facilitated by the forming of the inter-unit salt bridge interactions SB5 and SB6 (TM1(i)-ECL(i+1)) on the extracellular side. Subsequent loss of SB5 and SB6 interactions leads to the complete opening of the channel, which is facilitated by breaking of several classes of inter-unit salt bridge interactions spread all over the channel (*≈* 10 as shown in Table 6) as well as an inter-unit hydrogen bond (Y94-D16). This is also promoted by the loss of BB-BB H-Bonds, loosening the TM helices (Fig. lOD). Further, the non-polar amino acids L17 and A18 that sit between the labels on the intracellular side are reducing the repulsion between the positively charged labels and are restricting the extent of the opening of the channel.

## Conclusions

For the first time, the activation mechanism of engineered MscL was studied at atomic level using mixture of an all-atom un-biased equilibrium, non-equilibrium MD simulations, path optimizing SMwST and free energy estimation algorithms. In the equilibrium MD simulations, the labels facilitated opening of the engineered MscL channel, although not completely, which may be due to the time scale associated with the opening of the channel. This opening of the channel began near the labels, and the salt bridge interactions between the extracellular loops and the TM helices restricted the opening of the channel on the extracellular side, thereby restricting the complete opening. Non-equilibrium simulations were used to open the channel completely, and the resultant non-equilibrium configurations were optimized using the string method with swarms of trajectories. This was accompanied by free energy estimation along the optimized transformation pathway. To test the stability of the active state, an un-biased MD simulations on final frames of non-equilibrium simulations were conducted, which resulted in an open/active conformation of the engineered MscL channel. The open/active conformation captured in FE simulations is not as open as had been previously proposed. Overall, according to the results presented above, ECL loops aid in the activation of this engineered channel which plays a key role in the opening of the channel. Also, the loss of several BB-BB H-Bonds and several SC-SC salt bridge interactions facilitate the opening of the channel.

## Acknowledgement

This research is supported by National Science Foundation grant CHE 1945465 and the Arkansas Biosciences Institute. This research is part of the Blue Waters sustained-petascale computing project, which is supported by the National Science Foundation (awards OCI-0725070 and ACI-1238993) and the state of Illinois. This work also used the Extreme Science and Engineering Discovery Environment (allocation MCB150129), which is supported by National Science Foundation grant number ACI-1548562. This research is also supported by the Arkansas High Performance Computing Center, which is funded through multiple National Science Foundation grants and the Arkansas Economic Development Commission.

## Supporting Information Available

Figures S1–S7 and Tables S1-S2 in Supporting Information provide additional analysis based on our MD and string method trajectories as discussed in the manuscript.

