## Supporting Information for "Elucidating the Molecular Basis of pH Activation of an Engineered Mechanosensitive Channel"

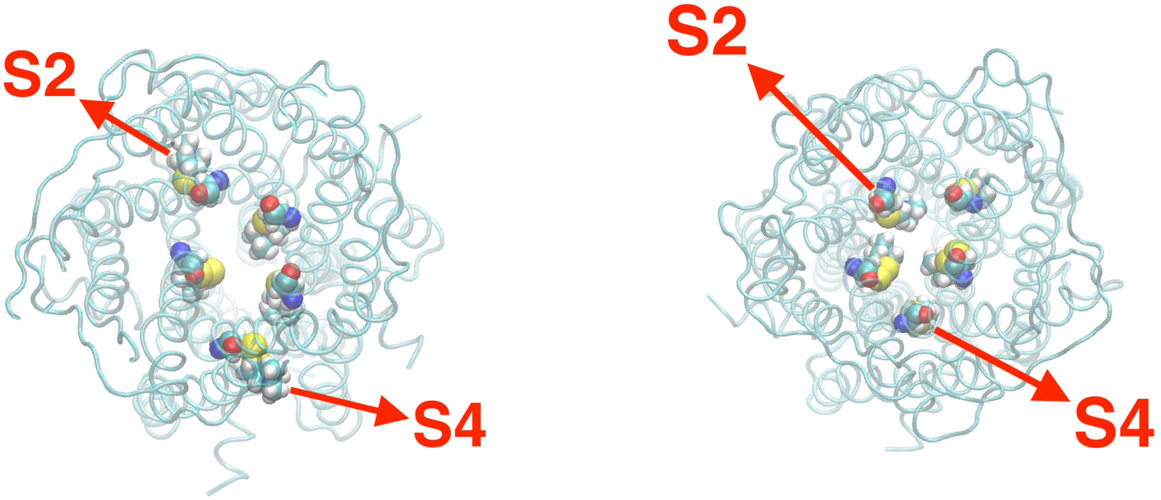


**Fig. S1.** Engineered MscL channels. Left is the engineered MscL (5-MTSET) with labels attached to all the 5 domains, and the labels attached to 2&4 oriented perpendicular to the normal and other 3 oriented parallel. Right is same as the left except that the labels attached to 2&4 are adjusted similar to the other 3(5-MTSET(A)).

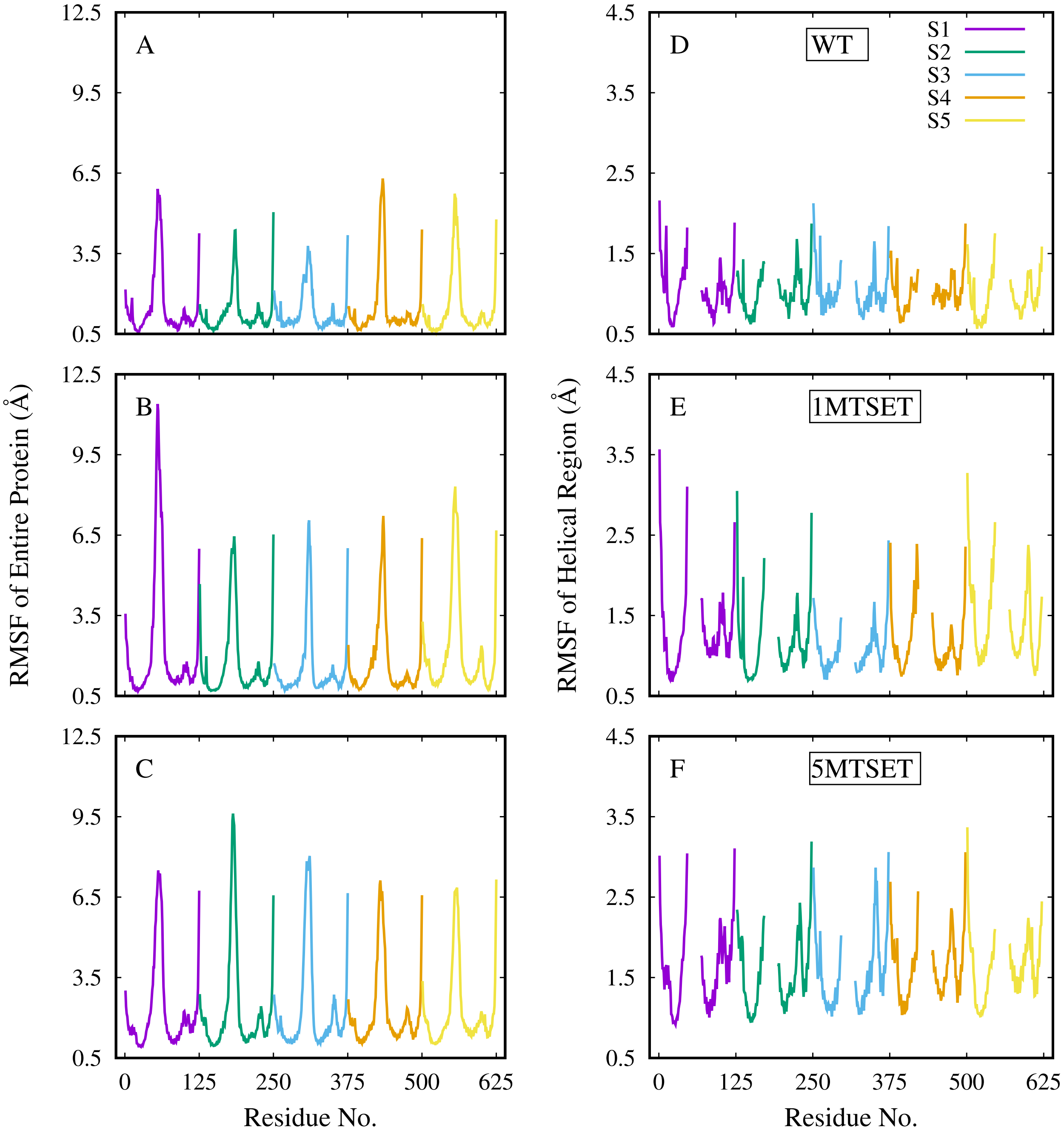


**Fig. S2.** RMSF vs. residue number. (A-C) Reflect the RMSF of entire protein. (D-F) Reflect the RMSF of just TM helices. (A, D) are WT, (B, E) 1-MTSET, and (C, F) 5-MTSET. Entire trajectory was used to calculate the RMSF.

**Table S1: Top 10 residues interacting with labels attached to chain-A.**

| System | Interacting Residues | | | | | | | | | |
|---|---|---|---|---|---|---|---|---|---|---|
| | **V21** | **V19** | **L17** | **D16** | **G24** | **N13** | **I14** | **I23** | **A18** | **V15** |
| **1-MTSET** | 100% | 100% | 100% | 100% | 100% | 98% | 28% | 15% | 10% | 0.0% |
| **5-MTSET** | 100% | 100% | 100% | 99% | 79% | 86% | 99% | 79% | 72% | 23% |

**Table S2: Residues interacting with labels sorted according to interaction frequency.**

| System | Interacting Residues | | | | | | | | |
|---|---|---|---|---|---|---|---|---|---|
| 1-MTSET | **V21** | **V19** | **L17** | **N13** | **D16** | **G24** | | | |
| | 100% | 100% | 100% | 100% | 98% | 98% | | | |
| 5-MTSET | **V21** | **V19** | **L17** | **I14** | **D16** | **G24** | **N13** | **I23** | **A18** |
| | 100% | 100% | 100% | 99% | 99% | 86% | 79% | 79% | 72% |

Residues only with $>60\%$ interaction frequency are considered. In the case of 5-MTSET more residues interact with the labels compared to the 1-MTSET(Table S2). All the top 10 residues that are interacting with the labels belongs to TM1 and they are all non-polar except N13 and D16 (Table S1).

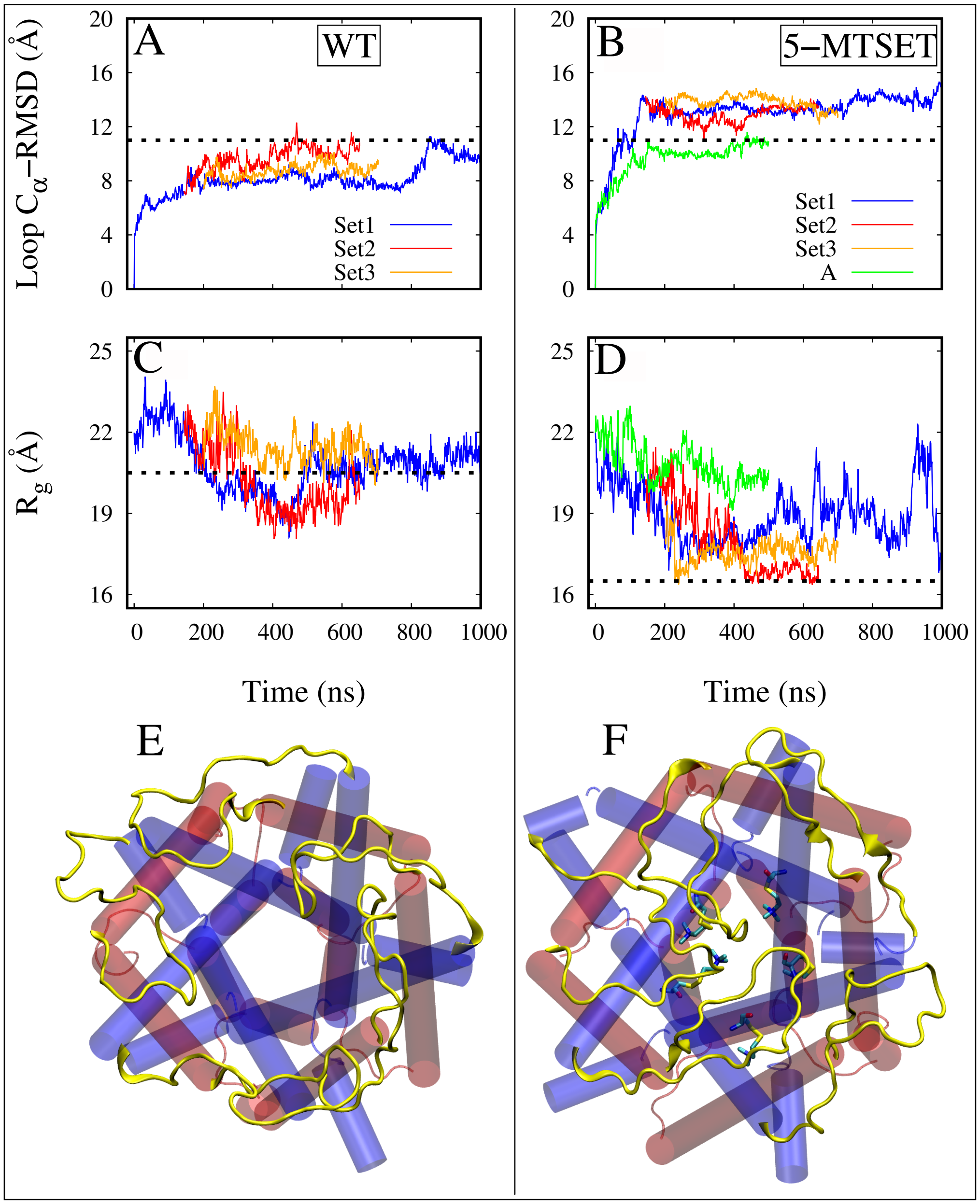


**Fig. S3.** RMSD ($C_\alpha$) and radius of gyration ($R_g$) of extracellular loops as a function of time. (A-B) RMSD's of all WT systems (A) were below 11 Å whereas in the engineered systems they were all greater than 11 Å. (C-D) $R_g$ of WT systems was stabilized around 20.5 Å, where as the longest simulations in the case of 1-MTSET and 5-MTSET systems were reduced to 16.5 Å. (E-F) Representative snap shots of WT and 5-MTSET(*) systems highlighting the loops behavior.

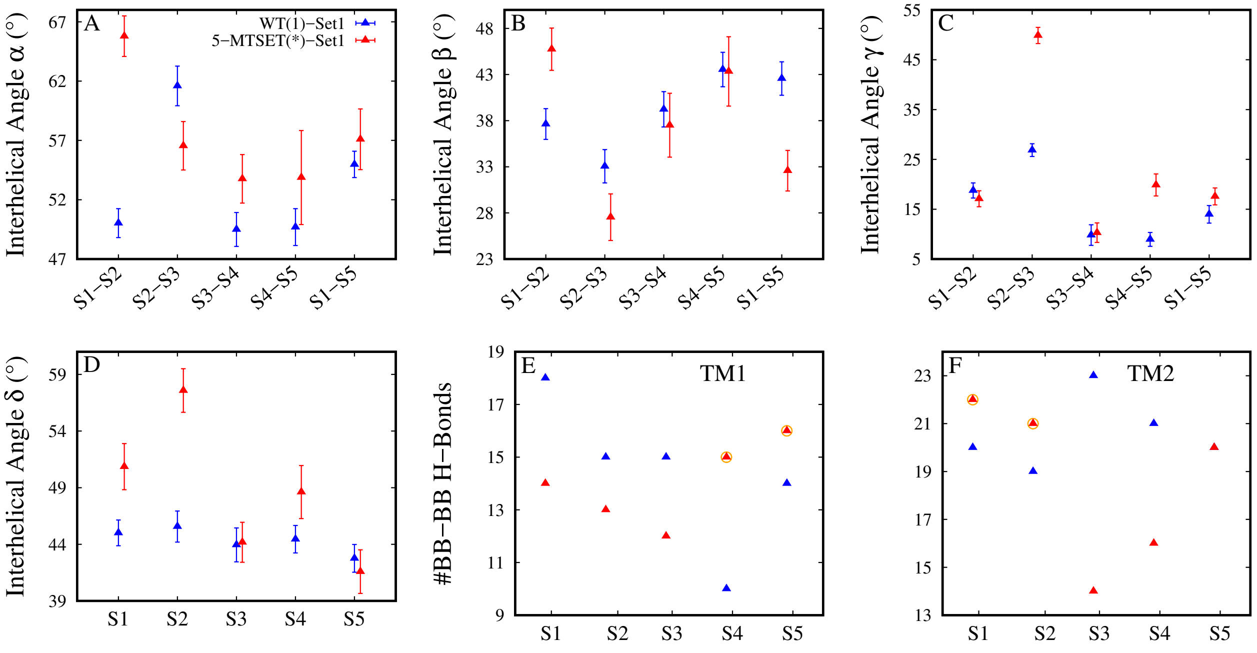


**Fig. S4.** Orientation angles and backbone-backbone hydrogen bonds of open MscL structure. (A-D) Orientation angles $\alpha$, $\beta$, $\gamma$, and $\delta$. (E-F) Number of backbone-backbone hydrogen bonds of TM1 and TM2 helices respectively.

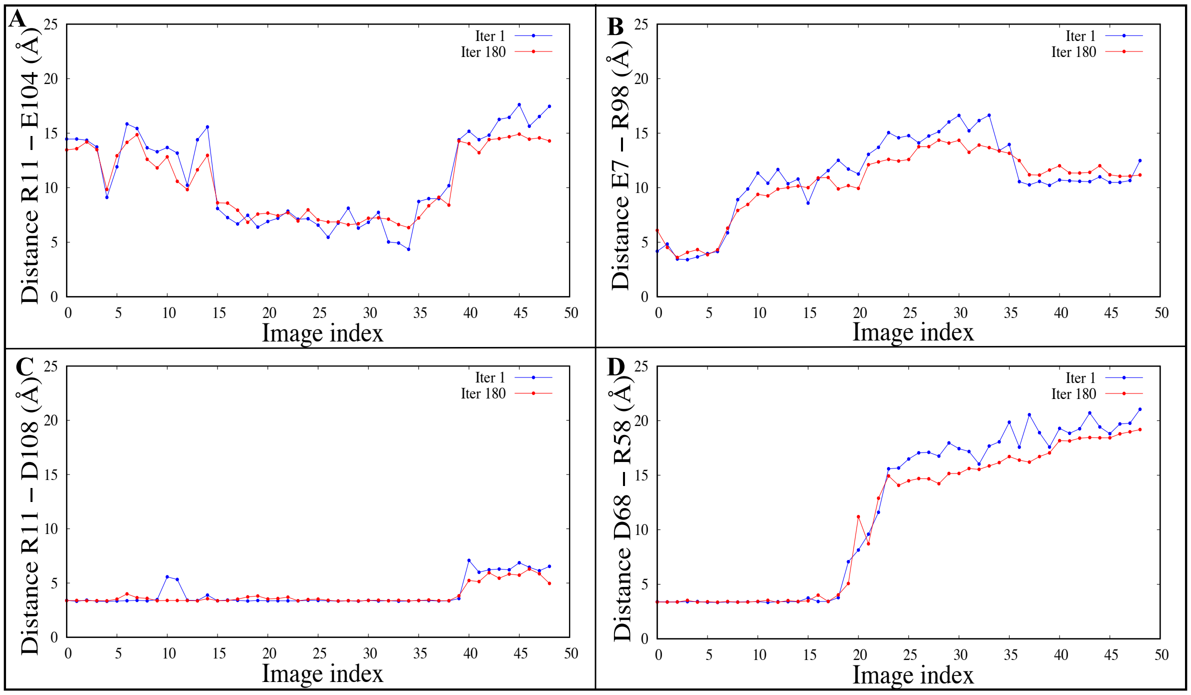


**Fig. S5.** Inter and intra unit salt bridge involved in the activation of the protein . (A-D) salt bridge distance between TM1 & TM2 R11-E104(SB1)(A), salt bridge distance between TM1 & TM3 E7-R98(SB2)(B), salt bridge distance between TM1 & TM3 R11-D108(SB3)(C), salt bridge distance between ECL1 & ECL1 D68-R58(SB3)(D)

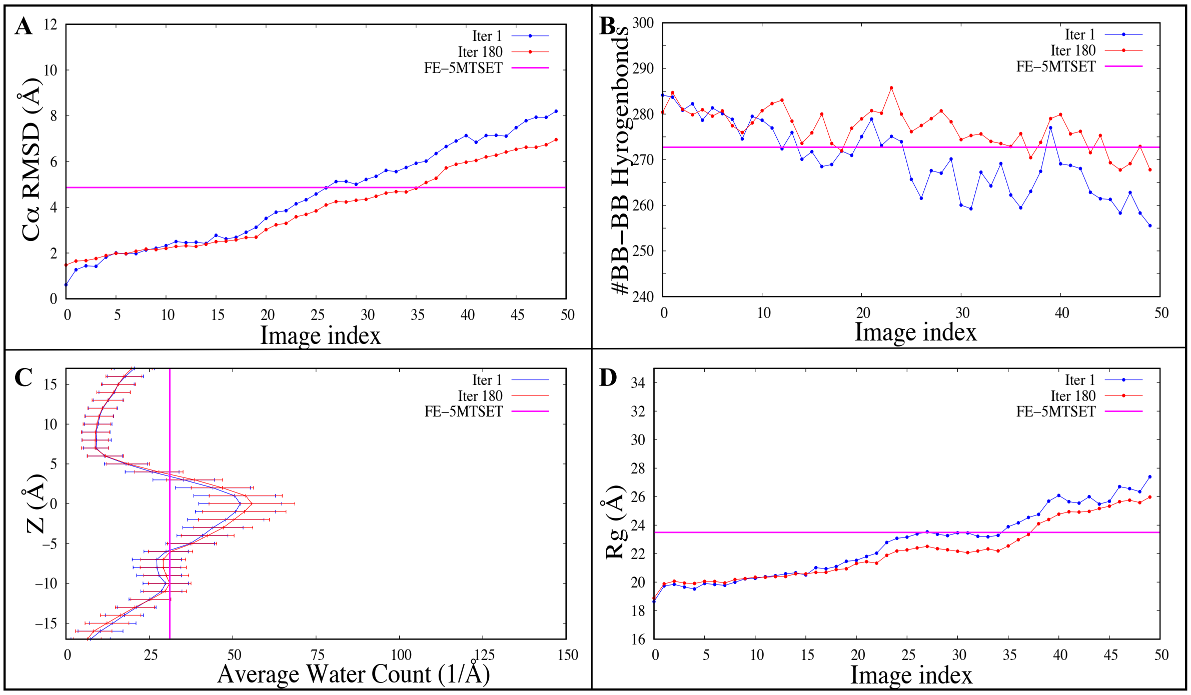


**Fig. S6.** RMSD, # BB-BB hydrogen bonds, average water count and radius of gyration($R_g$). (A-D) protein C$\alpha$ RMSD (A), # BB-BB H-bonds (B), average water count across the pore (C) and radius of gyration ($R_g$) between Iterations 1 , Iterations 180 systems, and FE NE simulation of 5MTSET . Iterations 1 , Iterations 180, and FE NE 5MTSET data are represented in blue, red and magenta respectively. The complete data of iteration 1 and 180 are considered for the above calculation each point value is averaged over 20 copies of each image. last 100ns simulation data was considered for the average FE NE simulation values.

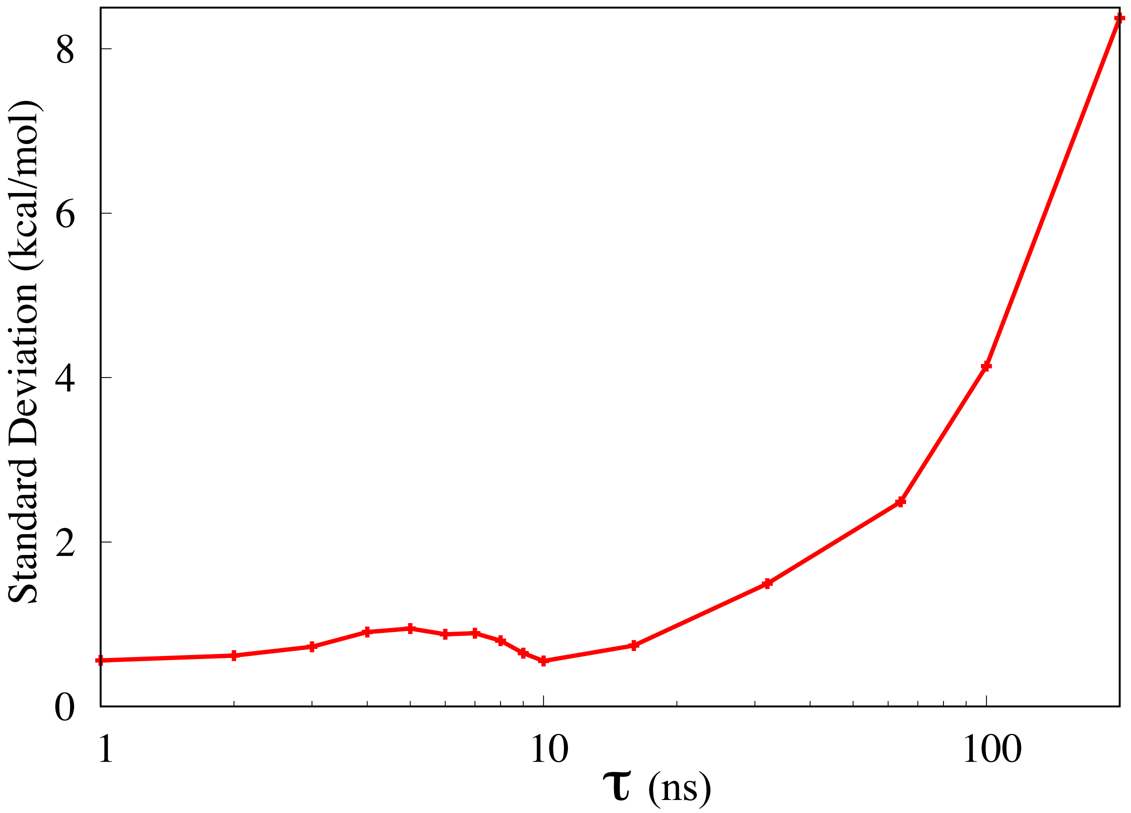


**Fig. S7.** Standard deviation chart of free energies calculated for each $\tau$ values.
